# Orderly mitosis shapes interphase genome architecture

**DOI:** 10.1101/2025.06.03.657645

**Authors:** Krishnendu Guin, Adib Keikhosravi, Raj Chari, Gianluca Pegoraro, Tom Misteli

## Abstract

Genomes assume a complex 3D architecture in the interphase cell nucleus. Yet the molecular mechanisms that determine global genome architecture are only poorly understood. To identify mechanisms of higher order genome organization, we performed high-throughput imaging-based CRISPR knockout screens targeting 1064 genes encoding nuclear proteins in multiple human cell lines. We assessed changes in the distribution of centromeres at single cell resolution as surrogate markers for global genome organization. The screens revealed multiple major regulators of spatial distribution of centromeres including components of the nucleolus, kinetochore, cohesins, condensins, and the nuclear pore complex. Alterations in centromere distribution required progression through the cell cycle and acute depletion of mitotic factors with distinct functions altered centromere distribution in the subsequent interphase. These results identify molecular determinants of spatial centromere organization, and they show that orderly progression through mitosis shapes interphase genome architecture.

## Introduction

Genomes are complex polymers. In the cell nucleus, the genome is organized via several structures at different length-scales. At the shortest scale, genomic DNA is wrapped around a histone octamer to form a nucleosome (Wen, Fang et al. 2025). Strings of nucleosomes then fold onto themselves to form a chromatin fiber, which in turn organizes itself into chromatin loops, typically in the 1-100 kb range. Chromatin fibers further fold into 0.5-2 Mb topologically associated domains (TADs), which can homotypically associate with each other into transcriptionally active A compartments and transcriptionally repressed B compartments spanning several megabases (Rowley and Corces 2018, Hildebrand and Dekker 2020, Misteli 2020, Paldi and Cavalli 2025). While these genome features occur in most cell types and species, all chromatin features exhibit extensive single-cell variability (Finn and Misteli 2019).

The organization of genomes is non-random within the cell nucleus (Parada and Misteli 2002, Oliver and Misteli 2005, Bouwman, Crosetto et al. 2022). Chromosomes and individual gene loci tend to occupy preferred positions relative to the nuclear boundary and relative to each other (Shachar and Misteli 2017, Scholz, Sumida et al. 2019, Bouwman, Crosetto et al. 2022). For example, the chromosomes that contain clusters of ribosomal genes congregate in 3D space to form the subnuclear compartment of the nucleolus (Dundr, Misteli et al. 2000). Similarly, transcriptionally repressive genome regions are often associated with the nuclear lamina at the periphery of the cell nucleus and around the nucleolus (Croft, Bridger et al. 1999, Alagna, Thomas et al. 2023). Defects in spatial genome organization are associated with multiple diseases including cancer and accelerated aging (Misteli 2010, Wang, Luo et al. 2023, Amodeo, Eyler et al. 2025).

Recent studies have shed light on to the mechanisms that determine the local organization of the genome (Bonev and Cavalli 2016, Finn and Misteli 2019, Misteli 2020). Chromatin loops and domains are formed by loop extrusion, in which the ringlike condensin protein complex acts as a molecular motor (Terakawa, Bisht et al. 2017) to extrude the chromatin fiber to form loops, which can eventually congregate into TADs (Ganji, Shaltiel et al. 2018, Davidson and Peters 2021). In contrast, the molecular mechanisms that determine the higher-order global genome organization, such as the location of genes, chromatin domains or chromosomes within the 3D space of the nucleus, are less clear. Some insights come from the observation in yeast and *C. elegans* that transcriptionally repressive chromosomes are preferentially tethered to the nuclear periphery via histone modifications (Towbin, Gonzalez-Aguilera et al. 2012, Shachar and Misteli 2017). Furthermore, unbiased screening approaches have suggested that progression through S-phase is essential to establish the nuclear location of individual genes (Joyce, Williams et al. 2012, Shachar, Voss et al. 2015). It has also been suggested that the propensity to undergo homotypic interactions promotes the clustering of similar genome regions, for example, the association of ribosomal genes in the nucleolus (Lafontaine, Riback et al. 2021) or the formation of intranuclear heterochromatin blocks (Misteli 2020).

The centromere is a prominent structural feature of all chromosomes (Murray and Szostak 1985). Centromeres are specialized genomic loci that assemble the kinetochore protein complex, which connects chromosomes with the microtubule spindle during mitosis, and through their attachment ensure error-free chromosome segregation (McKinley and Cheeseman 2016). Like other chromosomal features, centromeres have been observed to assume non-random locations in the cell nucleus across species. In yeast, centromeres cluster and localize at the nuclear periphery at some or all stages of the cell cycle (Guin, Sreekumar et al. 2020). Variable degrees of clustering have been observed in apicomplexan parasites (Bunnik, Venkat et al. 2019), plants (Fransz, De Jong et al. 2002), flies (Padeken, Mendiburo et al. 2013), and mice (Weierich, Brero et al. 2003, Stevens, Lando et al. 2017), where peri-centromeres cluster into prominent chromocenters, presumably via homotypic interactions (Brandle, Fruhbauer et al. 2022). In humans, centromere clustering is less pronounced but increased clustering of centromeres near nucleoli has been observed in multiple cell lines (Weierich, Brero et al. 2003, Bury, Moodie et al. 2020, Rodrigues, MacQuarrie et al. 2023, Kumar, Gholamalamdari et al. 2024), particularly prominently in human stem cells where most centromeres are localized near nucleoli (Wiblin, Cui et al. 2005, Rodrigues, MacQuarrie et al. 2023). The fact that clustered centromeres tend to dissociate from the nucleolus during stem cell differentiation (Rodrigues, MacQuarrie et al. 2023) may point to a functional role of nucleolar centromere clustering. However, the underlying molecular mechanisms determining spatial centromere distribution remain elusive.

Given their prominent nature and non-random location in the cell nucleus, we used centromeres as proxies for higher order spatial genome organization to identify molecular determinants of global genome architecture. We tested 1064 chromatin-associated proteins in high-throughput imaging (HTI)-based CRISPR/Cas9 knockout (KO) screens in human cell lines to identify conserved molecular determinants of nuclear centromere distribution. Our data identifies proteins implicated in diverse biological functions. Furthermore, by impairing the function of several of these candidates during the cell cycle, we demonstrate that defective mitotic progression alters centromere distribution in the daughter cells. We conclude that orderly progression through mitosis shapes global 3D genome architecture.

## Results

### The spatial distribution of centromeres is cell-type specific

We first sought to quantitatively profile the spatial distribution of centromeres in human cells (**Fig. 1**). We used HTI to visualize endogenous centromeres in eight human cell lines from different tissues and with distinct proliferation properties, including immortalized retinal pigment epithelium RPE1 cells, immortalized human HFF fibroblasts, the induced pluripotent stem cell (iPSC) line WTC-11, and several cancer cell lines of different origin (**Table S1**). Some of these cell lines contain numerical aberrations of chromosomes (Giard, Aaronson et al. 1973, Kotecki, Reddy et al. 1999), which were taken into account when setting the baseline for the quantitative analysis of centromere distribution in individual cell lines. Centromeres were visualized by indirect immunofluorescence for the integral kinetochore component CENP-C, which localizes to centromeres at all stages of the cell cycle and completely colocalized throughout the cell cycle with the centromere protein CENP-A (Hori, Amano et al. 2008, Klare, Weir et al. 2015). (**Fig. S1a**). CENP-C was used as marker for centromeres in all subsequent experiments.

**Fig. 1:**
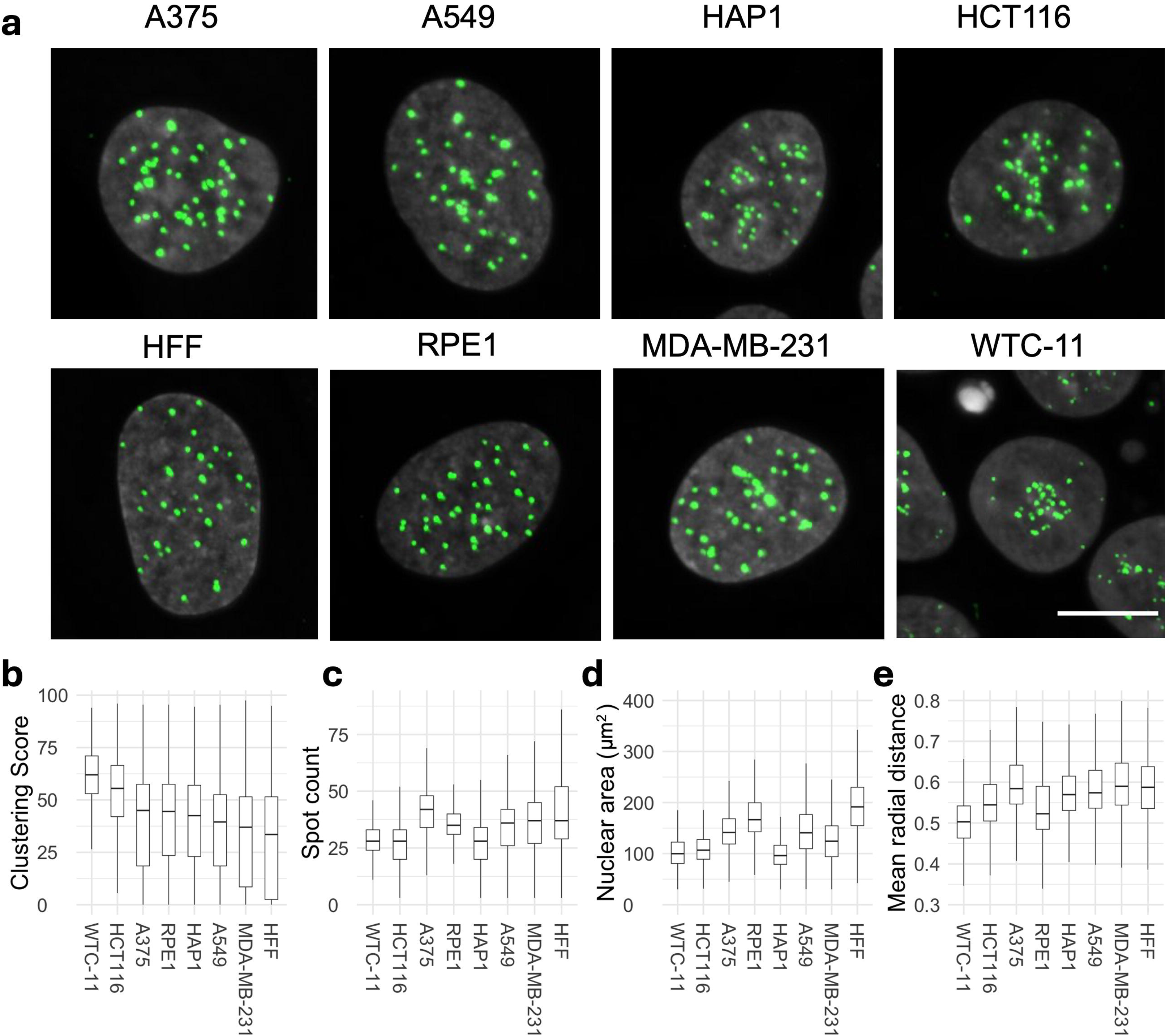
Spatial organization of centromeres is cell-type specific in human cell lines. **a**, Representative images of CENP-C (green) and DAPI (gray) stained nuclei in indicated human cell lines. Scale bar: 10 µm. **b**, **c**, Spatial organization of centromere quantified using Ripley K’s clustering score (**b**), CENP-C spot count (**c**). **d**, Nuclear area and **e**, mean radial distance in human cell lines. Statistical significance of difference between cell lines for clustering score, spot count, mean radial distance and nuclear area was tested using ANOVA (p-value or ‘Pr(>F)’ < 2e-16) following Tukey’s HSD test to compare means of all pairs of cell lines. Box plots represent the inter-quantile range (IQR) between first and third quantile (box), the median (horizontal bar), and the whiskers extend to the highest or lowest data point up to 1.5 times of IQR. Values are from one representative experiment with at least 7 technical replicates. At least 1000 cells were analyzed in each category per experiment.

We quantified the number of centromere spots per nucleus and analyzed centromere spatial distribution in the nucleus by HTI in several thousand cells per cell line by using HiTIPS, an open-source HTI analysis platform that accurately segments nuclei and centromeres in large HTI image datasets **(Fig. S1b**) (Keikhosravi, Almansour et al. 2024). For each cell, we measured the number of centromeres per nucleus as spot count and also derived a centromere clustering score, which measures the overall distribution of centromeres in the nucleus (**Fig S1c-d**, also see Methods). The clustering score is a metric derived from the Ripley’s K function, which we established in pilot experiments as a robust and sensitive measure of centromere clustering (Keikhosravi, Guin et al. 2025). The clustering score quantifies deviations of the centromere distribution from uniformly distributed spots, also known as complete state of randomness (CSR), and it is normalized to nuclear size. Importantly, the clustering score is robust to changes in the centromere spot number and thus accounts for any differences in centromere numbers in the various cell lines or due to aneuploidy (Keikhosravi, Guin et al. 2025).

We observed significant qualitative differences in centromere distribution amongst human cell lines (**Fig. 1a**). For example, in WTC-11 cells centromeres were strongly clustered, in line with centromere association with the nucleolus observed in other human stem cells (Wiblin, Cui et al. 2005, Rodrigues, MacQuarrie et al. 2023). In contrast, A549 basal epithelial cells derived from lung cancer and MDA-MB-231 epithelial-like breast cancer cells showed noticeably less clustering than HFFs, which exhibited the most dispersed distribution among the cell lines tested (**Fig. 1a**). These visual trends were confirmed by quantitative HTI analysis using the clustering score, which was highest for WTC-11 cells and lowest for HFFs (**Fig. 1b**). These differences in clustering were unrelated to spot number or to nuclear area (**Fig. 1c and 1d**). In addition, WTC-11 cells had the lowest and HFF cells the highest median population values for the mean normalized radial CENP-C distance, which represents the per-cell average distance of centromeres from the center of the nucleus (**Fig. 1e**), consistent with the differential clustering behavior in these two cell lines. Statistical analysis of variance (ANOVA) indicated that most cell lines were significantly different from each other based on centromere clustering score or spot count (**Table S2**), indicating cell-type specificity of centromere distribution patterns. We also noted cell-to-cell variation for all centromere distribution parameters within the population (**Fig. 1b, c, e**), demonstrating single-cell heterogeneity of centromere distributions as previously observed for various other features of genome organization (Finn and Misteli 2019). In conclusion, quantitative HTI analysis of centromeres localizations in thousands of single cells shows that spatial patterns of centromeres in the human cell nucleus are cell-type specific.

### Imaging-based CRISPR knockout screens identify regulators of centromere clustering

Having established the heterogenous and non-random nature of centromere clustering in the nucleus, we sought to identify the molecular basis for this phenomenon. To do so, we developed an arrayed HTI-based CRISPR-KO screening assay to identify regulators of the spatial distribution of centromeres (**Fig. 2a**). For the screens, we designed an sgRNA library targeting 1064 genes encoding nuclear proteins, enriched in structural components of the nucleus, epigenetic modifiers, and components of the genome maintenance and expression machinery (for library composition see **Table S3**). A non-targeting, scrambled sgRNA and sgRNAs targeting the non-expressing *OR10A5* gene were used as negative controls for sgRNA transfection and for CRISPR-induced DNA damage response (Liu, Golji et al. 2019), respectively. In addition, as a positive control for sgRNAs transfection, we used sgRNAs against the essential *PLK1* gene whose ablation results in rapid and extensive cell death (Schibler, Jevtic et al. 2023). As a positive control, we used sgRNAs targeting the condensin II complex component *NCAPH2,* whose silencing has previously been shown to induce clustering of centromeres (Hoencamp, Dudchenko et al. 2021). In light of our observation that spatial patterns of centromere distribution can be different between cell lines, we performed screens in two cell lines, HCT116 and RPE1, which represent clustered and unclustered centromere patterns, respectively (see **Fig. 1a**). For quantitative HTI analysis, we performed imaging-based phenotypic scoring of centromere distribution patterns using centromere spot count and the Ripley’s K-based clustering score as read-out parameters (Keikhosravi, Guin et al. 2025). All CRISPR-KO screens were performed in biological duplicates and generated data from a few hundred to over a thousand cells per replicate for each target gene (**Table S4 and S5**).

**Fig. 2:**
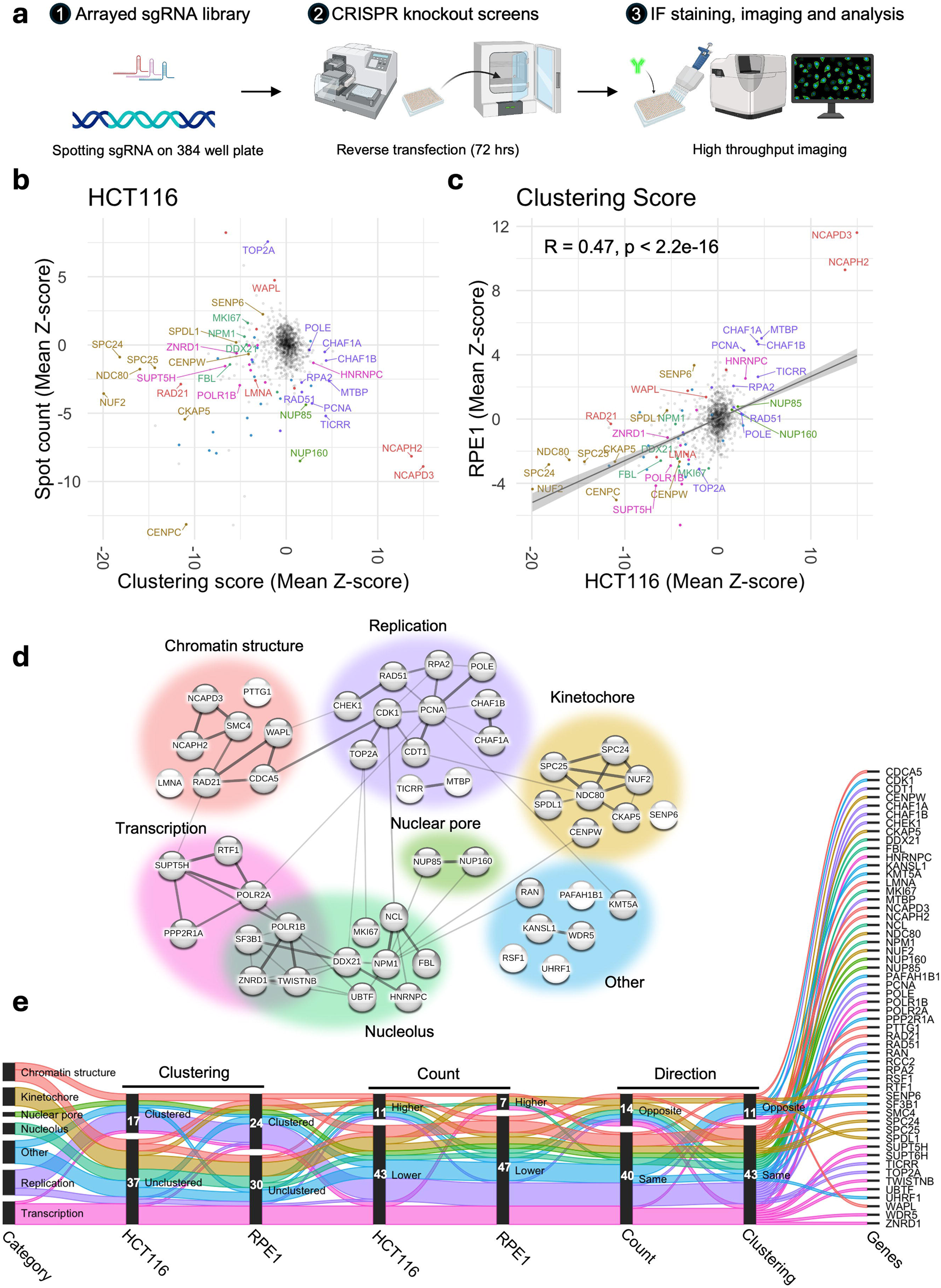
Identification of the molecular determinants of spatial centromere distribution across cell types. **a**, Schematics showing three stages of high-throughput imaging based arrayed CRISPR knockout screen employed to identify molecular determinants of spatial centromere distribution. **b**, Changes in spot count (mean Z-score of two replicates, y-axis) and clustering score (mean Z-score of two replicates, x-axis) for each of the 1064 sgRNAs. The most prominent hits were labelled and color coded as in Fig. 2d. Non-hits are colored in gray. **c**, Changes in clustering score in HCT116 (mean Z-score of two replicates, x-axis) and in RPE1 (mean Z-score of two replicates, y-axis) cells for each of the 1064 sgRNAs. Hits and non-hits are color coded and labelled as in **b**. A linear trend line (gray) was fitted to the data and Pearson’s correlation coefficient calculated is indicated at the top left corner of the plot. **d**, Network diagram with lines between 52 common hits drawn based on known physical and/or genetic interactions generated by the STRING database. Thickness of the lines indicates higher strength of data supporting the interaction. Broad categories are color coded as indicated. **e**, Plot shows changes in clustering (clustered or unclustered), count (higher or lower) and direction between two cell lines (same or opposite) for each of the common genes that are color coded based on their category as in Fig. 2d. Counts of genes in each subcategory are indicated. Values represent two biological replicates. Typically, 200-500 cells were analyzed for each target gene per experiment.

The results of the screens indicated consistent phenotypic separation of the positive and negative controls (**Fig. S2a, b**) and high reproducibility of hits in the two biological replicates for both cell lines (**Fig. S2c, d**). We defined hits as sgRNAs perturbations that altered either spot count or clustering by a Z-score of at least 2.5 units from the median phenotype of all sgRNAs included in the library (**Table S3**). Any sgRNA KO’s that resulted in a large number of dismorphic and/or abnormally sized nuclei, as measured by nucleus compactness and/or nuclear area, were filtered out from subsequent steps of the analysis. We excluded from analysis sgRNAs which resulted in high cytotoxicity (cell number Z-score < −2.5), or which produced inconsistent results across the two biological replicates (see Methods).

Following these criteria, we identified 111 genes whose CRISPR-KO altered centromere distribution in HCT116 cells (**Fig. 2b, Table S4**). Among these, 80% (89/111) altered the CENP-C clustering score, 41% (45/111) altered CENP-C spot count, and 20% (23/111) altered both parameters. The majority of hits (81%; 72/89) unclustered centromeres, whereas 19% (17/89) increased clustering (**Table S4**). Among the 23 genes that altered both parameters, six increased the clustering score and decreased spot count, indicating higher clustering, whereas the opposite trend was observed for four genes, indicating dispersion (**Table S4**). The remaining genes (13/23) concomitantly decreased the clustering score and spot count, suggesting global dispersion, but local clustering of centromeres into fewer but larger local clusters (**Table S4**). Concomitant increase of both spot count and clustering score was not observed. The effects on centromere distribution did not correlate with changes in nuclear area (**Fig. S2e, f**).

We similarly identified 113 hits when we performed the CRISPR-KO screen in RPE1 cells, which are characterized by a lower degree of centromere clustering than HCT116 cells (**Fig. S3, Table S5**). Similar to HCT116 cells, we observed a non-linear relationship of spot count and clustering score in RPE1 cells (**Table S5**). The majority (77%, 87/113) of identified sgRNAs altered centromere spot count, 40% altered the clustering score, while 17% altered both parameters (**Fig. 2c, Table S5**). When analyzed using the clustering score, the majority (58%, 26/45) of hits dispersed centromeres, whereas the rest (42%, 19/45) increased clustering (**Table S5**).

Select hits were orthogonally validated using siRNA knockdown with a validation rate of 90% (27/30) (**Fig. S4a**). Reassuringly, in line with known centromere-nucleoli association (Bury, Moodie et al. 2020), several nucleolar proteins, including NPM1, NCL and FBL, were identified as hits, confirming the validity of our screening approach. In addition, our positive control NCAPH2 represented in the library and another condensin II component NCAPD3 were strong hits in both cell lines and in all replicates of the screen. We identified both essential and non-essential genes and only very few hits altered cell cycle distribution, indicating that the hits were not due to secondary effects on the cell cycle or cell viability (**Fig. S4b-c**). In addition, centromere clustering levels were generally similar in G1, S and G2/M phases for most hits compared to scrambled control except for a handful of cases where the pattern changed upon knockdown of target genes (**Fig. S4d**).

A comparative analysis of the CRISPR-KO screen results in HCT116 and in RPE1 cells indicated that knockout of most genes similarly altered centromere distributions in both cell lines, but that the extent of change (Z-score) could vary depending on the initial state of centromere distribution (R = 0.47, p < 10^-10^, **Fig. 2c**). We identified 52 genes that alter centromere distribution in both cell lines (**Fig. S5a**). The majority of these genes altered phenotypes in the same direction for clustering score (79%, 41/52) and spot count (73%, 38/52) (**Fig. 2e**). Only rare examples of cell-type specific opposite effects were observed (**Fig. 2e and S5b**). Similarly, we identified genes that altered centromere distribution in only one cell line (**Fig S5b, Table S4 and S5**). Taken together, these data identify both conserved and cell-type specific regulators of centromere distribution.

To gain insights into the functions of the common hits we used STRING analysis which identifies pathways based on known physical and genetic interactions (**Fig. 2d**) (Szklarczyk, Nastou et al. 2025). Based on this analysis, centromere distribution modifiers were grouped into six categories: regulators of chromatin structure, kinetochore proteins, nucleolar proteins, nuclear pore complex components, replication factors and transcription-associated factors (**Fig. 2d**). Interestingly, while knockout of most replication- and nuclear pore- associated genes increased clustering, knockout of kinetochore components and transcription-associated factors led predominantly to centromere dispersion in both cell types (**Fig. 2e and S5b**). Loss of proteins implicated in chromatin structure or the nucleolus either clustered or dispersed centromeres in a gene-specific manner (**Fig. 2e and Fig. S5b**). It is also important to note that factors grouped into these broad classes may perform functions in multiple categories. For example, the NUP107-160 complex which is a prominent structural component of the nuclear pore is known to also interact with kinetochores (Orjalo, Arnaoutov et al. 2006). These results identify major regulators of spatial centromere organization.

### Spatial re-distribution of centromeres requires cell-cycle progression

The identified regulators of centromere distribution are involved in diverse cellular functions and pathways, suggesting multi-layered control of centromere distribution. To gain mechanistic insight, we asked whether the identified regulators act at particular points in the cell cycle. To establish a baseline for analysis, we quantitated centromere distribution in G1, S and G2/M of cell-cycle staged HCT116 and RPE1 cells based on DAPI and EdU pulse-labelling as described before (Salic and Mitchison 2008, Bruhn, Kroll et al. 2014, Roukos, Pegoraro et al. 2015) (**Fig. 3a, b**). As expected, due to the duplication of the genome during replication, the number of detectable centromere spots increased in S phase cells (p = 0.009) and was highest in G2/M HCT116 cells (p< 10^-10^ **Fig. 3c).** A marginal increase in clustering score was observed in G2/M cells compared to G1 cells, (**Fig. 3d**; p = 0.002) while radial positioning of centromeres remained mostly unchanged except for a small increase in G1 cells (**Fig. 3e**; p = 0.004). A similar trend was observed in RPE1 cells for all three parameters (**Fig. 3c-e**). As expected, the nuclear area was significantly increased in S and G2/M in both HCT116 and RPE1 cells (**Fig. 3f**, p < 10^-10^). The lack of strong correlation between nuclear area and clustering score within G1, S or G2/M subpopulations (R < 0.3) indicates that increased clustering scores in G2/M cells are unrelated to nuclear size increase (**Fig. 3g**). We conclude that, in line with observations on radial position of genomic loci (Shachar, Voss et al. 2015) and of chromosome territories (Jowhar, Gudla et al. 2018), the overall distribution of centromeres does not vary strongly within the interphase of the cell cycle.

**Fig. 3:**
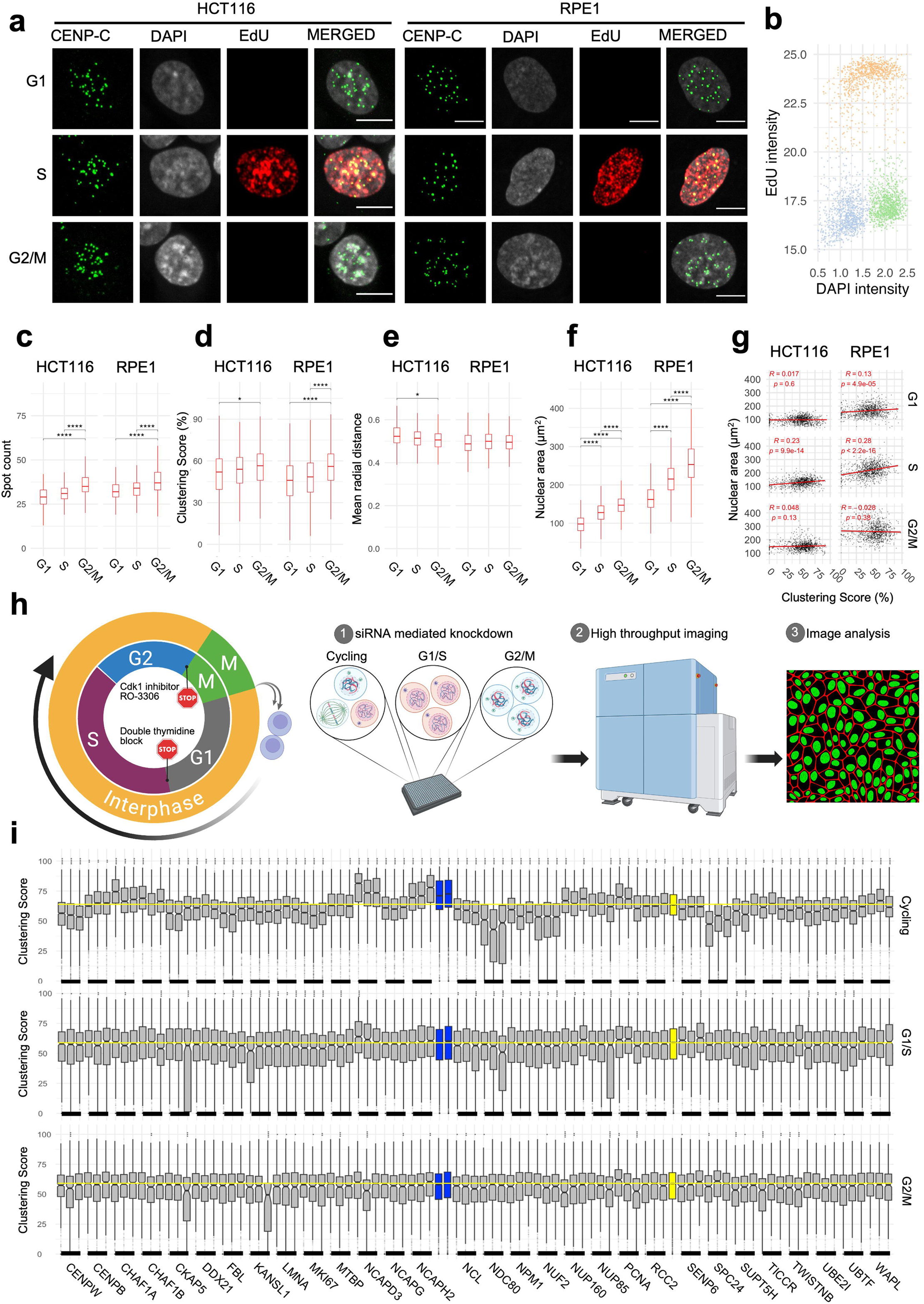
Changes in spatial organization of centromeres require progression through the cell cycle. **a**, HCT116 and RPE1 cells at G1, S and G2/M phases stained with CENP-C (green), DAPI (gray) and EdU (red). Scale bar: 10 µm. **b**, EdU intensity (y-axis) and DAPI intensity (x-axis) showing separation between G1 (brown), S (gray), and G2/M (green) sub populations in cycling HCT116 cells. Comparison of cells at G1, S or G2/M stages (x-axis) for their clustering score (c), spot count (**d**), mean radial distance (**e**), or nuclear area (**f**). Statistical significance of differences was tested by pairwise t-test with Bonferroni correction. Asterisks indicate level of significance between a given pair reflecting the corresponding p-value of that comparison. **g**, A linear regression line (red) fitted through the single cell data for nuclear area (y-axis) and clustering score (x-axis) in cells at different cell cycle phases in HCT116 and RPE1 cells. Pearson correlation coefficient and respective adjusted p-values are indicated at the top of each panel. **h,** Experimental outline to test cell cycle stage-specific effect of knocking down select hits. **i**, Effect of siRNA knockdown for a panel of genes (x-axis) using three individual siRNAs per gene in HCT116 cells that are either arrested at G/S and G2 or cycling. Two control siRNAs for siNCAPH2 are in blue and siScrambled in yellow. Mean value for siScrambled is depicted by a horizontal yellow dotted line. Statistical significance of differences was tested by performing pairwise t-tests with Bonferroni correction using siScrambled as control group. Box plots represent the inter-quantile range (IQR) between first and third quantile (box), the median (horizontal bar), and the whiskers that extend till the highest or lowest value up to 1.5 times of IQR. Values are from one representative experiment. Typically, 200 to 500 cells were analyzed in each category. Statistical significance of difference was denoted by stars where * indicates p ≤ 0.05, ** indicates p ≤ 0.01, *** indicates p ≤ 0.001 and **** indicates p < 0.0001.

To specifically ask whether the identified hits required progression through the cell cycle, we performed siRNA knockdown of 30 select hits in asynchronous cells or in cells that were either arrested at the G1/S boundary by standard double thymidine block (Chen and Deng 2018) or at the G2/M boundary by treatment with the CDK1 inhibitor RO-3306 as previously described (Vassilev, Tovar et al. 2006) (**Fig. 3g**; see Methods).

Loss of cell viability upon transfection of siDeath, which simultaneously targets several essential genes, as compared to siScrambled (**Fig. S6a**) and reduction in NCAPH2 protein upon transfection of siNCAPH2 (**Fig. S6b**) indicate efficient siRNA knockdown in cycling, G1/S and G2/M cells. While centromere distribution was altered upon knockdown of these genes in cycling cells as expected, no changes in centromere distribution were observed when knockdowns were done in G1/S or G2/M arrested cells (**Fig. 3h).** We conclude that while the distribution of centromeres does not vary during the cell cycle, progression through the cell cycle is required to bring about changes in centromere distribution in the absence of key regulators of centromere clustering. These results demonstrate that the identified modifiers of centromere distribution do not act in the maintenance of centromere distribution during interphase.

### Normal progression through mitosis is required for faithful interphase centromere distribution

Having established that cell cycle progression is required for the effects of the identified centromere distribution factors, we asked at what stage of the cell cycle the centromere distribution factors act. We measured changes in clustering score before and after progressing through either S phase or mitosis in cells depleted of a given factor (**Fig. 4a**). We selected for this analysis four proteins which all interact with centromeres and contribute to efficient and error-free chromosome segregation during mitosis but all have distinct functions: NCAPH2 is a component of the Condensin II complex and responsible for axial compaction of chromosomes (Shintomi and Hirano 2011, Green, Kalitsis et al. 2012, Gibcus, Samejima et al. 2018); KI67 is a well-known marker of cell proliferation that decorates nucleoli in interphase cells and coats chromosomes during mitosis (Booth, Takagi et al. 2014, Cuylen, Blaukopf et al. 2016); SPC24 and NUF2 are kinetochore components and part of the NDC80 complex which connects the kinetochore to microtubules (McCleland, Kallio et al. 2004, Ciferri, Pasqualato et al. 2008). . Auxin-inducible degron cell lines to deplete NCAPH2 and KI67 have previously been characterized (Takagi, Ono et al. 2018). In addition, we generated dTAG-SPC24 and NUF2-dTAG cell lines by CRISPR knock-in into HCT116-Cas9 parental cells (**Fig. S7a-b**; Methods). Homozygous knock-in in select clones was verified by PCR genotyping (**Fig. S7c-d**) and correct localization and expression of dTAG-SPC24 and NUF2-dTAG as compared to the parental cell line was confirmed by indirect immunofluorescence staining of the tagged proteins and by Western blotting, respectively (**Fig. S8a-b, e-f**) (McCleland, Kallio et al. 2004). Effective depletion of each factor by more than 90% as assessed by western blotting was achieved within 3 hours (**Fig. S8c-d**).

**Fig. 4:**
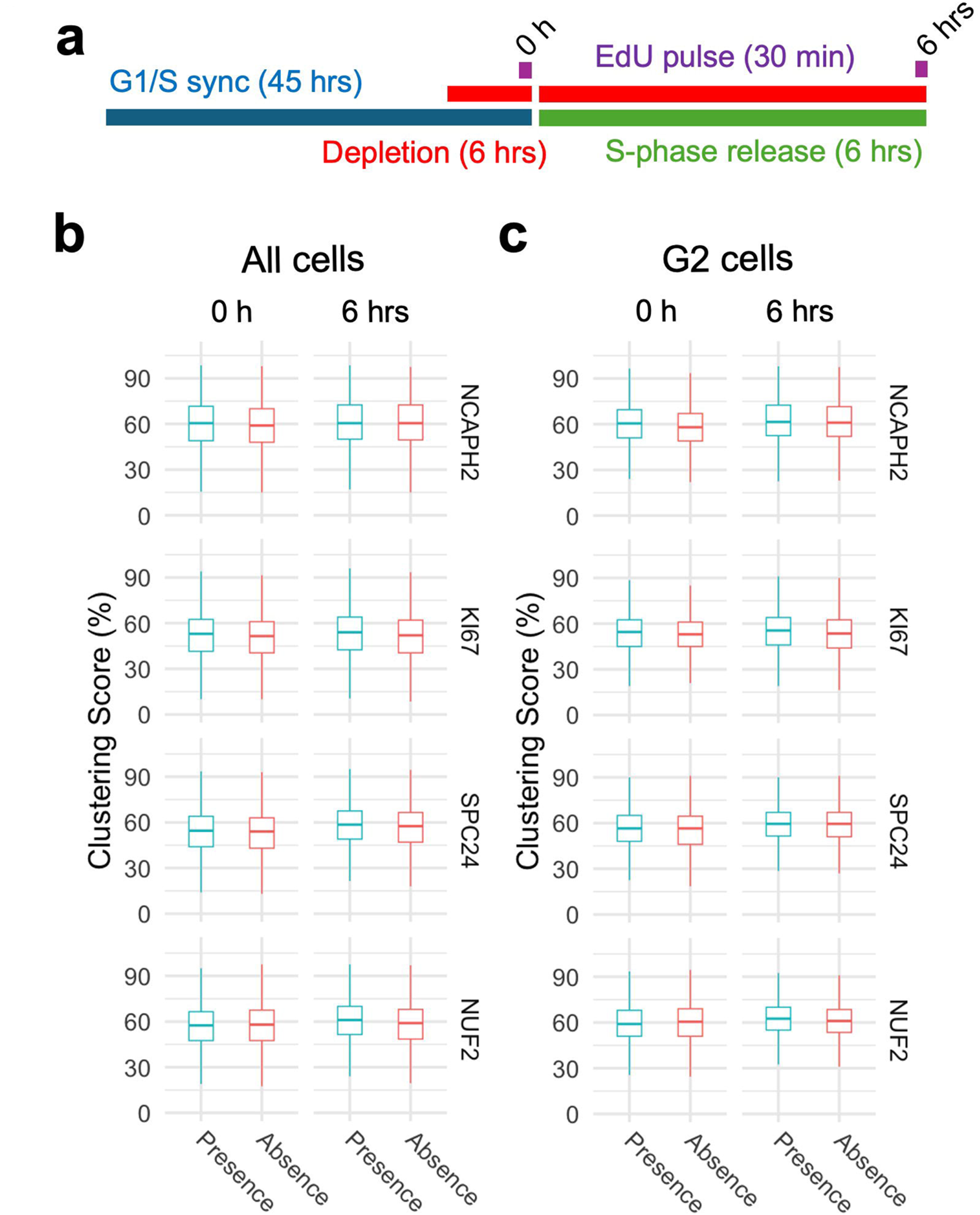
Progression through S-phase in the absence of select clustering factors does not alter interphase genome organization. **a**, Experimental outline to compare centromere distribution during progression through S-phase in the presence or absence of clustering factors. **b**, **c**, Clustering score in all cells (b) or G2/M cells (c) in the presence (blue) or absence (red) of indicated clustering factors before and after S-phase release from G1/S arrest. Pair-wise comparisons were tested with t-test with Bonferroni correction, and the level of significance is indicated by asterisks if any. Pairs without significant difference are not labelled. Box plots represent the inter-quantile range (IQR) between first and third quantile (box), the median (horizontal bar), and the whiskers that extend till the highest or lowest value up to 1.5 times of IQR. Values are from one representative experiment with three technical replicates. Typically, 200 to 500 cells were analyzed in each category.

Using these four HCT116-based degron lines, we acutely depleted individual factors specifically in cells arrested at the G1/S or G2/M boundaries and then released the cell cycle block (see Methods). We validated effective depletion of SPC24 and NUF2 in G1/S and G2/M arrested cells (**Fig. S9a-d**). First, we compared clustering scores in cells progressing through S-phase in the presence or absence of NCAPH2, KI67, SPC24, or NUF2 (**Fig. 4a**). Upon release from a standard double thymidine block, the majority (58-78%) of cells reached G2/M after 6 hours (**Fig. S10a and S10c**). Progression through S phase was equally efficient in the presence or absence of KI67, SPC24 or NUF2 (**Fig. S10a and S10c).** Acute NCAPH2 depletion mildly delayed S-phase progression (**Fig. S10a and S10c**), as reported earlier (Rodemoyer, Kariyawasam et al. 2025). Although loss of KI67 has been reported to delay replication of centromeres and pericentromeric loci (van Schaik, Manzo et al. 2022, Stamatiou, Huguet et al. 2024), no effect on bulk S-phase progression after Ki67 loss was observed in our hands (**Fig. S10a**). No effect on Clustering Scores was evident as cells progressed through S-phase into G2, regardless of the presence or absence of any of these proteins (p > 0.05; **Fig. 4b and c**). We conclude that these centromere distribution modifiers do not act in S-phase.

Next, we tested if the loss of function of these centromere distribution modifiers during mitosis altered centromere localization in the subsequent interphase cells (**Fig. 5a)**. HCT116 cells were arrested at the G2/M boundary by treatment for 20 hours with the CDK1 inhibitor RO-3306 as previously described (Vassilev, Tovar et al. 2006), and then released for 6 hours in the absence of each mitotic factor. The newly formed G1 cells were analyzed for centromere distribution. As expected, mitotic progression in the absence of SPC24 or NUF2 was slowed upon release from the G2/M block (McCleland, Kallio et al. 2004) with 28-35% of cells reaching G1 after 6 hours compared to 58-60% in the presence of SPC24 or NUF2 (**Fig. S10b and S10d)**. As expected, some aberrant nuclear phenotypes were evident in the absence of SPC24 or NUF2 (**Fig. S11a-d**). Mitotic progression in the absence of NCAPH2 and KI67 were similar to that of control cells (**Fig. S10b and S10d**). While the centromere distribution phenotypes remained unaltered compared to cycling cells in the presence of these proteins, progression through a single mitosis in the absence of any of these proteins altered centromere distribution phenotypes in the subsequent G1 phase as assessed by quantitation using the Ripley’s K clustering score (**Fig. 5b and c**) and visual inspection (**Fig. 5d**). Loss of NCAPH2 had the largest effect and resulted in increased clustering of centromeres in G1 cells (**Fig. 5c**; p <10^-10^). Similarly, progression through mitosis in the absence of KI67 reduced clustering in the new G1 cells (p = 1.27e-08), as did loss of SPC24 (p < 10^-10^) or NUF2 (p < 10^-10^) (**Fig. 5c**). We conclude that the function of NCAPH2, KI67, SPC24 and NUF2 during mitosis determines centromere distribution patterns in the newly formed daughter nuclei. The fact that loss of proteins with distinct mitotic functions perturb centromere organization in the subsequent G1 phase suggests that, rather than their specific mitotic functions, it is the orderly progression of cells through mitosis that is required to ensure the faithful maintenance of spatial centromere distribution.

**Fig. 5:**
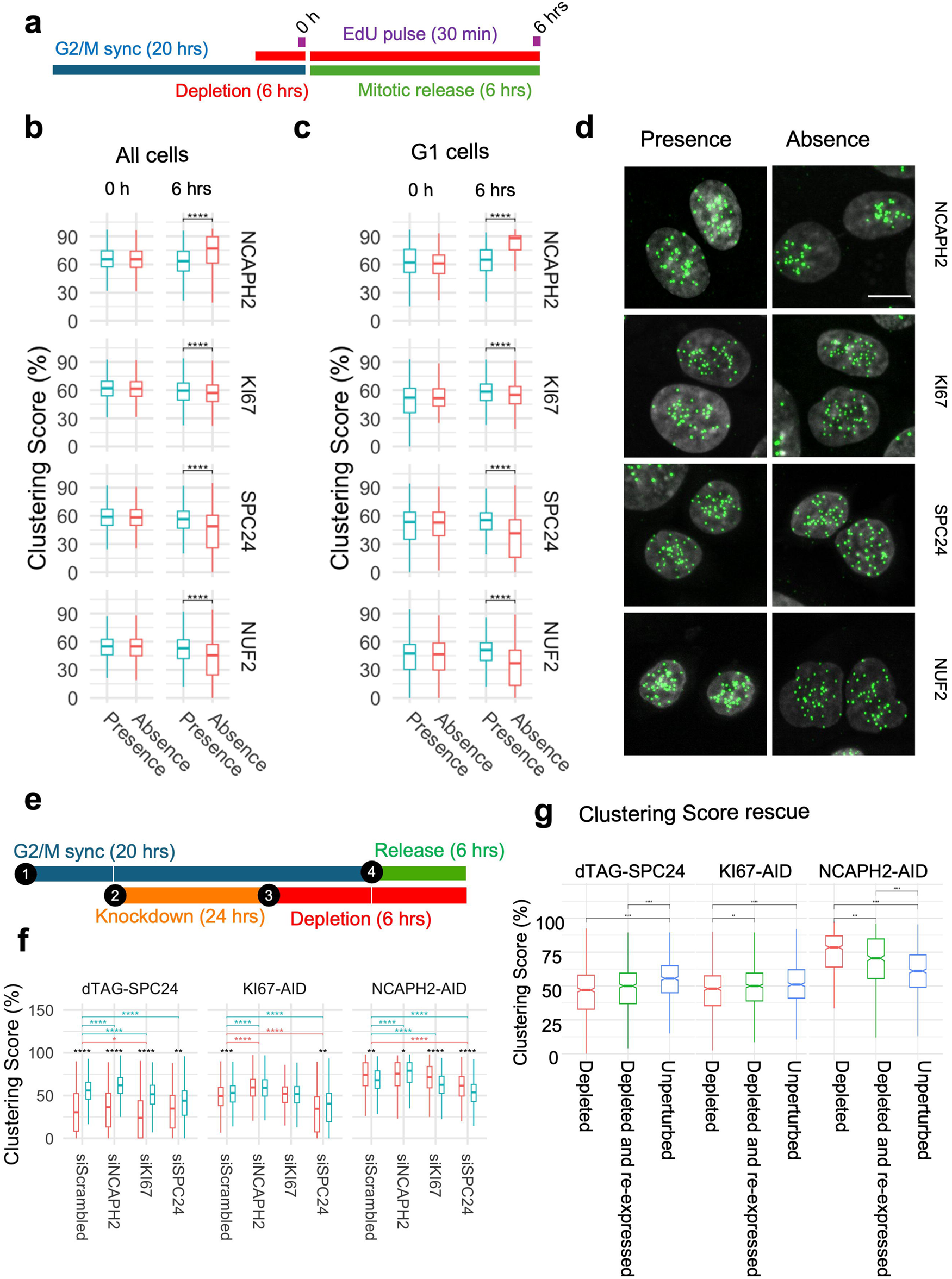
Orderly progression through mitosis is required to re-establish centromere distribution. **a**, Experimental outline to compare centromere distribution during mitotic progression in the presence or absence of clustering factors. **b**, **c**, Clustering score in all cells (b) or G1 cells (c) in the presence (blue) and absence (red) of indicated clustering factors before (0 h) and after (6 hrs) mitotic release from G2 arrest. Pair-wise comparisons were tested with t-test with Bonferroni correction, and the level of significance is indicated by asterisks. Pairs without significant differences are not labelled. **d**, Representative images showing G1 nuclei stained with DAPI (gray) and CENP-C (green) in the presence or absence of indicated factors. Scale bar: 10 µm. **e**, Schematics for co-depletion of indicated factors. **f**, Clustering score (y-axis) in G1 cells after siRNA knockdown of indicated factors (x-axis) in presence (blue) or absence (red) of SPC24, KI67 or NCAPH2 as indicated. Statistical significance of difference between indicated pairs were tested by performing t-test with Bonferroni corrections for multiple comparisons and denoted by stars, where * indicates p ≤ 0.05, ** indicates p ≤ 0.01, *** indicates p ≤ 0.001 and **** indicates p < 0.0001. Comparisons between individual siRNA groups (x-axis) and inter-group comparisons in the presence and absence of the given degron tagged protein are shown in black, blue (presence) and red (absence), respectively. **g**, Clustering score (y-axis) in cells that were depleted (red) or depleted of indicated factors and then re-expressed (green) or remained unperturbed (blue). Statistical significance of difference between indicated pairs were tested by performing t-test with Bonferroni corrections for multiple comparisons and denoted by stars, where higher number of stars indicate higher confidence levels. Box plots represent the inter-quantile range (IQR) between first and third quantile (box), the median (horizontal bar), and the whiskers that extend till the highest or lowest value up to 1.5 times of IQR. Values are from one representative experiment containing three technical replicates. Typically, 200 to 500 cells were analyzed for each category. Statistical significance of difference was denoted by stars where * indicates p ≤ 0.05, ** indicates p ≤ 0.01, *** indicates p ≤ 0.001 and **** indicates p < 0.0001.

Since all four factors act during mitosis but had different effects on centromere distribution, we explored co-depletion phenotypes to understand functional overlap between these factors, if any. We combined siRNA knockdown and degron-based depletion of NCAPH2, KI67 or SPC24 in pairwise combinations along with scrambled siRNAs and non-depleted cells as controls (**Fig. 5e**). We observed additive effects as simultaneous loss of SPC24 and KI67 further reduced the clustering score than the individual loss of either KI67 (p = 5.0640e-47) or SPC24 (p = 5.8800e-03). Similarly, the centromere unclustering upon KI67 knockdown was rescued by simultaneous NCAPH2 depletion (**Fig. 5f**; p= 3.2280e-10). In contrast, centromeres did not cluster more when *NCAPH2* was either knocked down (p= 5.9760e-56) or depleted (p= 3.2280e-10) in the absence *SPC24* (**Fig. 5f**) indicating that SPC24 functions upstream of NCAPH2 in regulating spatial centromere position. These findings point to an intricate interplay of these factors and pathways in determining centromere positioning.

We finally asked whether the aberrant altered centromere distribution in daughter cells upon depletion of mitotic factors can be reversed upon re-expression of NCAPH2, KI67 or SPC24. To test this idea, each of these factors were depleted for 6 hours in asynchronous cells following washout of degron ligands to allow re-expression of NCAPH2, KI67 or SPC24 as they progress through the cell cycle for 24 hours. Cells with and without depletion are used as controls. We observed partial rescue upon re-expression of all three factors as clustering scores partially returned towards that of the unperturbed cells (**Fig. 5g**).

Taken together these findings demonstrate a requirement for orderly progression through mitosis to shape the spatial distribution of centromeres and global genome organization in the subsequent interphase nuclei.

## Discussion

We identify here several cellular factors that determine the 3D positions of centromeres in the human cell nucleus, and we find that interference with orderly progression though mitosis alters centromere location in the subsequent interphase. We conclude that mitotic events shape the spatial organization of the interphase genome.

The most prominent group of centromere distribution effectors were components of the mitotic machinery, particularly multiple kinetochore proteins including all four components of the NDC80 complex (McCleland, Kallio et al. 2004) and components of the CENP-T-W-S-X complex (Nishino, Takeuchi et al. 2012). The fact that loss of multiple factors with distinct mechanisms of action, but all affecting mitosis, resulted in altered centromere distribution in the newly formed G1 cells points to a prominent role for orderly progression through mitosis as a main determinant of interphase centromere distribution, reminiscent of prior observations on lamina-associated chromatin domains which stochastically reposition during mitosis (Kind, Pagie et al. 2013).

A likely mechanism for the observed altered arrangement of centromeres in early G1 upon interference with mitotic machinery is the aberrant alignment of chromosomes in the mitotic plate and their uncoordinated migration towards the spindle poles (**Figure 6**). As cells enter mitosis, the outer kinetochores assemble on the centromeres, chromosomes condense and align on the metaphase plate. This process is initiated in late G2 when the KMN (KNL1, MIS12, and NDC80) complex, including the NDC80 complex, is loaded onto the kinetochore to stabilize microtubule attachments (Gascoigne and Cheeseman 2013). Loss of NDC80 components, such as SPC24 or NUF2, weakens microtubule attachments, but does not completely disrupt chromosome segregation as has been observed for CENP-A, CENP-C and other components of inner kinetochore (Ciferri, Musacchio et al. 2007). As such, chromosomes will progress through mitosis but will be imprecisely oriented in the metaphase plate and will migrate in an un-coordinated fashion to the spindle poles, leading to their dispersal in early G1. Indeed, we find that the assembly and disassembly of the NDC80 complex correlates with lower clustering score in G1 cells compared to G2/M cells in a cycling population. This effect is further exaggerated upon knockout or depletion of multiple NDC80 complex components resulting in stronger centromere dispersion. The observed mitotic effects on interphase organization are reminiscent of recent observations on the relationship of chromosome location and mis-segregation defects (Klaasen, Truong et al. 2022) where single-cell observations indicated that the more peripheral a chromosome is in the interphase nucleus, the higher its chance of improper alignment in the metaphase plate and consequently being mis-segregated leading to aneuploidy (Klaasen, Truong et al. 2022, Vukusic and Tolic 2022). Similarly, the observed effect of NCAPH2 depletion on centromere distribution may reflect a defect in chromosome segregation. Loss of NCAPH2 has been shown to lengthen chromosomes which may facilitate homotypic centromere-centromere interactions, resulting in the observed increase in clustering of centromeres in G1 (Hoencamp, Dudchenko et al. 2021). A further contributor to the mitotic effect on interphase centromere distribution may be defects in mitotic exit as suggested by our identification of KI67 as a determinant of centromere distribution. KI67 has been localized to centromeres (van Schaik, Manzo et al. 2022) and reported to act in late telophase as a surfactant to generate mechanical forces that are required for re-establishing nuclear-cytoplasmic compartmentalization in G1 cells (Cuylen, Blaukopf et al. 2016, Hernandez-Armendariz, Sorichetti et al. 2024). Loss of KI67 may disrupt the arrangement and progression of chromosomes in late telophase leading to redistribution of centromeres in G1. Pairwise depletion of these factors producing additive effects on clustering pointed to an intricate interplay of the affected pathways in determining interphase centromere positioning.

**Figure 6:**
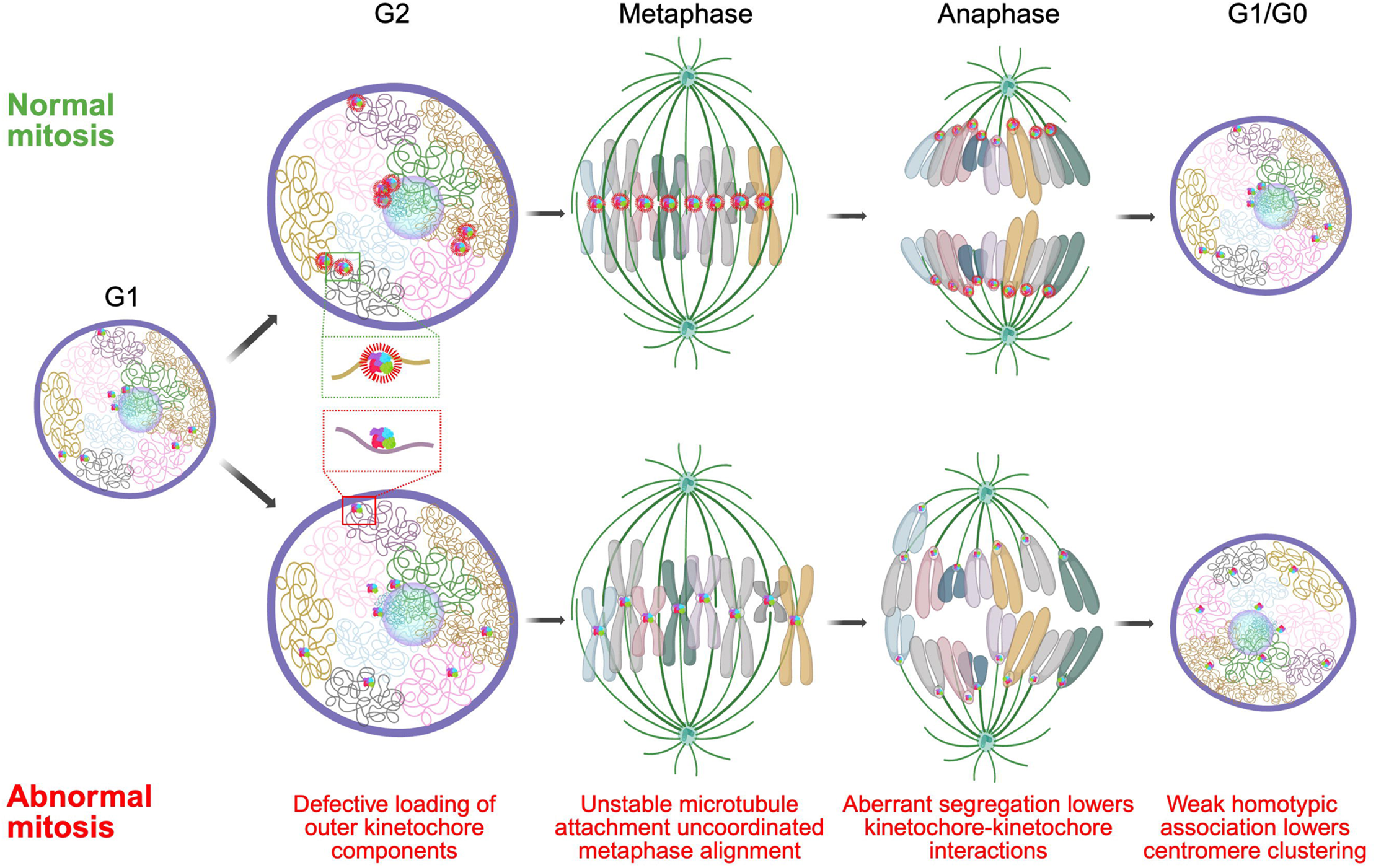
Mitotic events shape interphase genome organization. A model showing defective loading of outer kinetochore (inset; green and red box) in late G2 leads to uncoordinated metaphase alignment and aberrant migration towards spindle pole during anaphase that lowers chances of interactions between centromeres during telophase and the lack of homotypic adhesion results in dispersion of centromeres in the daughter nuclei.

Regardless of the precise molecular mechanisms for how mitotic progression affects the distribution of centromeres in interphase, it is likely that the location of centromeres in the nuclear space is in large parts driven by homotypic interactions. Centromeres are specialized genomic loci that are highly heterochromatic with low transcription activity (Altemose, Logsdon et al. 2022). It is well established that homotypic chromatin regions, such as heterochromatin, self-interact as is evident by the formation of the A and B chromatin compartments, which incorporate regions of similar chromatin status from distinct chromosomes (Hildebrand and Dekker 2020, Misteli 2020). A homotypic self-organization model for centromeres is in line with the presence of chromocenters in mouse, fly and plant cells, which represent clusters of peri-centromeric regions from multiple chromosomes forming large heterochromatin blocks (Sullivan and Karpen 2004, Probst and Almouzni 2011, Jagannathan, Cummings et al. 2019), but are largely absent in humans indicating factors implicated in chromocenter maintenance are probably not required for spatial centromere organization in humans. Indeed, we find that knockout of *HMGA1,* whose gene product stabilizes chromocenters in mouse (Jagannathan, Cummings et al. 2018), or other HMG genes, did not affect the spatial distribution of centromeres in human cells, suggesting species-specificity of some determinants of genome organization. A hetero-chromatin-driven homotypic interaction model also explains the prominent association of centromeres with the nucleolus in human stem cells, which are largely devoid of nuclear heterochromatin blocks (Meshorer and Misteli 2006), making the nucleolus the most prominent high-affinity binding site for centromeres in the nucleus. Our model is also in line with the long-standing observation that following mitosis, chromosome unfolding leads to the re-establishment of the chromatin landscape of interphase nuclei (Belmont and Bruce 1994).

It is intriguing to speculate that altered centromere position may have functional consequences. For example, dispersion of centromeres may increase centromere to non-centromere contacts thus influencing the local chromatin environment of both repositioned centromere and non-centromere loci possibly altering their transcriptional output. On the other hand, clustering of centromeres may help preserve integrity of the centromeric and peri-centromeric chromatin environment but may also increase the likelihood of inter-centromere translocations thereby disrupting genome stability. Finally, identification of centromere position regulators now allows experimental perturbation of higher-order genome organization and test its impact on overall gene expression and genomic stability, adding to our current understanding of the broad connection between genome structure and function.

## Methods

### Cell Culture

All cell lines used in this study are grown in a humidified 37°C incubator in presence of 5% CO_2._ Sources, composition of growth media, relevant references of the respective cell culture protocols are in Table S1. Cells were fixed with 2% paraformaldehyde (PFA, Electron Microscopy Sciences, Cat. No. 15710) solution in media by adding one equal volume of 4% PFA solution in PBS to the cell growth medium in each well for 15 minutes at room temperature. To activate protein degradation 500 nM auxin (Millipore Sigma, Cat. No. I3750-25G-A) or 1 µM dTAG13 (Tocris Bioscience, Cat. No. 6605) or dTAG^V-^1 (Tocris Bioscience, Cat. No. 6914) ligand was used.

### Cell cycle synchronization

Cells were synchronized at G1/S boundary by double thymidine block as described (Chen and Deng 2018). Briefly, cells were seeded at 20-30% confluency and grown for 24 hours following 18 hours of growth in media containing 2 mM thymidine (Sigma, Cat. No. T9250-5G). Cells were then washed with PBS and grown in fresh media for 9 hours. The growth media was changed with fresh media containing 2 mM thymidine. Cells were synchronized at the G2/M stage by treating cells with 9 µM RO-3306 (Millipore Sigma, Cat No. 217721-2MG) for 20 hours as described (Vassilev, Tovar et al. 2006). Cells were once washed in 300 µL fresh growth media and replenished with fresh growth media to release from the G1/S or G2/M block.

### Immunofluorescence staining

Indirect immunofluorescence (IF) staining was performed as previously described (Keikhosravi, Almansour et al. 2024, Keikhosravi, Guin et al. 2025). Briefly, fixed cells grown on a 96 well plate (Revvity, Cat. No. 6055300) or 384 well plate (CellVis, Cat. No. P384-1.5H-N) were washed with PBS and then permeabilized with 0.1% Triton X-100 (Sigma, Cat. No. T9284-500ML) solution in PBS for 15 minutes and again washed with PBS. These cells were blocked by incubating with 5% BSA (Millipore Sigma, Cat. No. A3294-100G) solution in PBST (0.05% Tween-20 in PBS) for 15 min at room temperature. Next, cells were incubated with appropriate primary antibody dilution prepared in blocking solution (5% BSA in PBST) for 1 hour at room temperature and then washed with PBS three times. Next cells were incubated for 1 hour with fluorescently labelled secondary antibody solution prepared in blocking solution at room temperature and then washed with PBS three times. DAPI (4′,6-diamidino-2-phenylindole) (Thermo Fisher Scientific, Cat. No. 62248) staining was performed by adding 5 µg/mL DAPI solution prepared in 1x PBS to the wells. Anti-CENP-C (MBL Biosciences, Cat. No. PD030) and anti-CENP-A (Abcam, Cat. No. AB13939) antibodies were diluted 1:1000 in blocking buffer (5% BSA solution prepared in PBST) and used for IF staining. In experiments where cells were EdU labelled, CENP-C primary antibody was directly conjugated with fluorophores using Mix-n-Stain™ CF® Dye Antibody Labeling Kit (Biotium, Cat. No. 922235). Anti-FLAG monoclonal antibody (Sigma, cat. No. F3165250) was diluted 1:250 in blocking buffer and used for IF staining.

### CRISPR-KO Library design

A custom arrayed synthetic sgRNA library targeting 1064 genes associated with chromatin biology and nuclear architecture was sourced from Synthego (Cat # SO17105 and 8311960), delivered lyophilized in 96-well plates, resuspended in RNAse free ddH_2_O, and reformatted in 384-well plate format using the PerkinElmer Janus and the Beckman Coulter ECHO525 liquid handlers at a final of 0.25 pmoles/µL. Each gene was targeted in the same well by 3 pooled sgRNA oligos that included the Synthego modified EZ Scaffold. The list of genes and sgRNA targeting sequences in the library is included in Table S3.

### CRISPR knockout screens

For reverse transfection of sgRNA oligos in 384-well format, 325 nL of library sgRNA 0.25 pmoles/µL were spotted in each empty well (0.08 pmoles/well) of an imaging plate (CellVis, Cat. No. P384-1.5H-N) using an ECHO525 acoustic liquid handler. As controls, and in each plate, we also spotted 7 wells each of non-targeting scrambled control sgRNA (Synthego, Cat. No. 063-1010-000-000), sgRNAs targeting each of *PLK1, OR10A5*, and *NCAPH2.* Three sgRNAs pooled together in gene knockout kits to target *PLK1*, *OR10A5* and *NCAPH2* were obtained from Synthego (Cat. No. GKO-HS1-000-0-1.5n-0-0). The control sgRNAs had the same chemistry and were spotted in the same quantities as the as the sgRNAs in the library. Spotted plates were dried at RT under a laminar flow cell culture hood, sealed, and stored at −30°C until the day of the reverse transfection.

The day of the transfection, the spotted sgRNA imaging plates were thawed and equilibrated at RT and then spun at 1400 rpm. The seal was removed, and 20 µL of prewarmed serum-free Optimem media (Thermo Fisher Scientific, Cat. No. 31985070)) was dispensed into each well for the imaging plate using a Thermo Fisher Multidrop dispenser. The ECHO525 was then used to dispense required amount of Lipofectamine RNAi MAX (Thermo Fisher Scientific, Cat. No. 13778075). The plates were then incubated at room temperature for 30 minutes to allow RNA-lipofectamine complexes to form. Next, a cell suspension prepared in prewarmed Optimem media (Thermo Fisher Scientific, Cat. No. 31985070) containing 20% FBS was dispensed into each well using the Multidrop for a total volume of 40 µl and an effective final sgRNA concentration of 2 nM. Plates were incubated at room temperature inside a laminar airflow hood for 30 minutes before they were transferred into cell culture incubator and allowed to grow at for 72 hours. CRISPR-KO screens were performed each in 2 biological replicates on different days.

### Imaging

384 well plates containing fixed cells were then stained for CENP-C and DAPI and imaged using a Yokogawa CV8000 spinning disk confocal microscope. Imaging parameters were as described before (Keikhosravi, Almansour et al. 2024). Briefly, Immunofluorescence images were collected on a multi-laser platform equipped with 405 nm (DAPI), 488 nm (for green fluorophores), 561 nm (for red fluorophores), and 640 nm (for far-red fluorophores) excitation lines that were combined through a 405/488/561/640 nm quad-band dichroic. Fluorescence was captured through a 60× water-immersion objective (NA = 1.2) and routed to either a 445/45 nm band-pass filter for DAPI or a 525/50 nm band-pass filter for green fluorophores, or 600/37 nm band-pass filter for red fluorophores or 676/29 nm band-pass filter for far-red fluorophores. A 16-bit sCMOS detector (2048 × 2048 pixels, 1 × 1 binning; effective pixel size = 0.108 µm) recorded Z-stacks with 1 µm steps, while real-time maximal projection was applied. Variable number of fields ranging from 9-22 were imaged per well in different imaging experiments to acquire sufficient number of cell images.

### Image analysis

Image analysis was performed as described (Keikhosravi, Guin et al. 2025). Raw image stacks were processed with HiTIPS, our previously described high-content analysis pipeline for fixed- and live-cell assay (Keikhosravi, Almansour et al. 2024). Max-projected DAPI channels provided nuclear masks, whereas CENP-A or CENP-C projections served for centromere spot localization (see **Fig. S1B**). Analysis settings in HiTIPS were adjusted to the typical nuclear diameter and the intensity/size characteristics of centromere foci. Nuclear segmentation employed the GPU-accelerated CellPose algorithm (Pachitariu and Stringer 2022), and centromere detection used a Laplacian-of-Gaussian approach. Final spot coordinates were defined as the centroid of each segmented focus. Average nuclear fluorescence intensity for the DAPI (405 nm) and EdU (640 nm) channels were measured at the single cell level. Clustering scores were calculated using a metric derived from Ripley’s K function as described (Keikhosravi, Guin et al. 2025).

### Identification of screen hits

Single cell data obtained from HiTIPS (Nucleus area, Number of CENP-C Spots per Cell, Ripley’s K clustering score, mean normalized CENP-C Spot Radial distance) were averaged on a per well level. Per well average values and nuclei counts was then used as an input for the screen statistical analysis using R 4.3.3 [(R Core Team (2024). _R: A Language and Environment for Statistical Computing_. R Foundation for Statistical Computing, Vienna, Austria. <https://www.R-project.org/>] and the cellHTS2 package (Boutros, Bras et al. 2006). Briefly, per well raw measurement were normalized on a per plate basis using the median value of the sgRNA library treatments and the B-score method. All the per plate normalized values in different library plates originated from a single biological replicate were further standardized using a robust version of the Z-score. The Z-score values for the same well and plate combination in different biological replicates were then averaged to obtain a Mean Z-score.

Screen hits were identified as genes whose knockout resulted in a mean Z-score either higher than 2.5 or lower than −2.5 for either Number of CENP-C Spots per Cell or Ripley’s K clustering score. Nuclei that were either abnormally shaped (solidity < 0.85) or were abnormally small (area < 30 µm^2^) indicative of micronuclei were not used for analysis. We also excluded sgRNAs which resulted in high cytotoxicity (Cell Number Z-score < −2.5) or whose Mean Z-score was smaller than the Z-score standard deviation of two replicates.

### DAPI and EdU labelling for cell cycle profiling

EdU labelling was performed using a kit (Thermo Fisher Scientific, Cat. No. C10340) as per manufacturer’s instructions. Briefly, cells were incubated with 1 ug/mL EdU for 45 minutes before fixation. Fixed cells were permeabilized and blocked as described for IF staining protocol. Next, genome incorporated EdU molecules during replication were fluorescently labelled by performing a click chemistry reaction for 30 minutes at room temperature. Subsequently cells were washed twice with 1% BSA solution in PBST and stained with 5 µg/mL of DAPI solution in PBS at room temperature for 1 hour.

### Image analysis for cell cycle profiling

Total nuclear fluorescence of the DAPI and EdU channel was quantified using HiTIPS (Keikhosravi, Almansour et al. 2024). The quantitation data was processed downstream using R packages and log2 transformed DAPI integrated intensity was used to distinguish the G1 and G2/M subpopulations. Similarly, EdU integrated fluorescence intensity was log2 transformed and cells with detectable EdU intensity as S-phase cells. Cells with DAPI intensity either higher than G2 cells (>4N population) or lower than G1 cells (subG1 population) were not analyzed.

### Construction of dTAG-SPC24 and NUF2-dTAG cell lines

Candidate guide RNAs targeting the N-terminus of SPC24 and C-terminus of NUF2 were designed using *sgRNA Scorer 2.0* (Chari, Yeo et al. 2017) and CRISPRor (Concordet and Haeussler 2018) (**Table S6**). Briefly, candidate guide RNAs were *in vitro* transcribed (IVT) and tested for cutting activity in cells, using an approach previously described (Gooden, Evans et al. 2021). Based on both the highest indel frequency as well as proximity to the point of insertion, candidate 207 was selected for SPC24 and candidates 286 and 289 were selected for NUF2. Oligonucleotides corresponding to these three guide RNAs were phosphorylated, annealed and ligated into either the pX458 backbone (Addgene #48138) (Ran, Hsu et al. 2013) for SPC24 (pJT142) or pDG458 (Addgene #100900) (Adikusuma, Pfitzner et al. 2017) for NUF2 (pMG1040). pSpCas9(BB)-2A-GFP (PX458) was a gift from Feng Zhang (Addgene #48138). Plasmid pDG458 was a gift from Paul Thomas (Addgene #100900).

To generate the homology directed repair (HDR) donors for construction of dTAG-SPC24 and NUF2-dTAG cell lines, first, DNA fragments with 5’ and 3’ homology arms were synthesized using Twist Biosciences and subsequently cloned into the pGMC00018 using isothermal assembly (Gibson, Young et al. 2009) to generate intermediate constructs pJT147 (SPC24) and pMG1063 (NUF2). Subsequently, the Puromycin-2A-dTAG-3X-FLAG and 3X-FLAG-dTAG-2A-Puromycin cassettes were then PCR amplified from an existing plasmid (contains 3X-FLAG tagged version of dTAG, generated from addgene #91796 or #91793) and then cloned into pJT147 to generate pJT152 and pMG1063 to generate pMG1064, respectively, using isothermal assembly approach. List of oligoes used to generate these constructs are listed (**Table S6**). All plasmids were sequenced completely using nanopore sequencing.

To generate dTAG-SPC24 homozygous knock-in line HCT116 Cas9 cells were co-transfected with two constructs: plasmid pJT142 encoding sgRNAs targeting the SPC24 locus and Cas9, and pJT152 encoding a donor template to introduce dTAG-FLAG and puromycin resistance using Lipofectamine LTX with Plus reagent (Thermo Fisher Scientific, Cat. No. 15338100) as instructed by manufacturer. Similarly, to generate NUF2-dTAG homozygous knock-in line HCT116 cells were co-transfected with pMG1040 encoding sgRNAs targeting NUF2 locus and Cas9, and pMG1064 encoding a donor template to introduce FLAG-dTAG and puromycin resistance. 24 hours after transfection, cells were washed with fresh media and grown in fresh media for 24 hours. Next, cells were selected for puromycin resistance by growing them in the presence of 1.5 µg/mL puromycin (Thermo Fisher Scientific, Cat. No. A1113803) for 72 hours. Every 24 hours, old media was replaced with fresh media containing puromycin. Cells were further expanded in presence of puromycin. Depending on the efficiency of transfection cells took 5-7 days to achieve confluence. The cells were then harvested by using trypsin (Gibco, Cat. No. 15050065). A portion of the cells were frozen to generate a polyclonal stock, and the remaining cells were taken forward for single cell cloning.

### Single cell cloning

To generate single cell clones, 500 cells were plated on a 15 cm dish and allowed to form colonies in the presence of puromycin. Single colonies appeared in 7-10 days. To isolate single colonies, 20-30 single colonies were selected, and a cloning cylinder was placed around them. The bottom of the cloning cylinder was sealed with grease. For clone isolation, 20 µL trypsin was added to each cylinder and cells from individual colonies were resuspended in separate wells in a 96 well plate to recover single cell clones.

### PCR confirmation

Genomic DNA from single cell clones was isolated using the Biorad reagent (Thermo Fisher Scientific, Cat. No. K0512) as recommended. To identify homozygously tagged dTAG-SPC24 clones KG49 (5’AGCTCAGACTTACAGGCGTG3’) and KG76 (5’TGATGGTGCTGATGGTTGCA3’) primers were used to amplify the genomic region flanking the site of integration at the SPC24 locus. This primer pair is expected to amplify a 2925 bp fragment from the tagged allele and an 1821 bp fragment from the untagged allele (Fig. S7a, and c). Similarly, homozygously tagged NUF2-dTAG clones were identified by PCR analysis using primer pair KG37 (5’CTGCTTTTCTTCCCCCACTG3’) and KG64 (5’AGAGGCAGCCTTTTCTCTGA3’). NUF2-dTAG alleles produced a 3046 bp amplicon while the untagged allele produced an amplicon of 1943 bp length (Fig. S7b, and d).

### Western blotting

Two PCR-confirmed single cell clones were used for western blot analysis to validate depletion of dTAG-SPC24 or NUF2-dTAG. Cells were grown in the presence of 1 µM dTAG13 ligand for 3, 6 and 9 hours. Cells grown in the absence of the ligand were grown as control. Protein samples were prepared using Bio-Rad lysis buffer (Bio-Rad, Cat. No. 1610747) as per the manufacturer’s instructions. Briefly, 10^6^ cells were collected, washed with PBS and lysed in 100 µL lysis buffer (Bio-Rad, Cat. No. 1610747) to isolate proteins.

Anti-FLAG monoclonal antibody (Sigma, Cat. No. F3165) was diluted 1:10000 in blocking buffer (5% BSA solution prepared in TBST) was used for detection of dTAG-SPC24 and NUF2-dTAG in western blots. For loading control anti-beta-tubulin antibody (Millipore Sigma, Cat. No. A2228) was diluted 1:25000 in blocking buffer and used in western blots. For comparison between untagged SPC24 and FLAG-dTAG-SPC24 protein levels antibody against SPC24 (gift from Todd Stukenberg lab) (McCleland, Kallio et al. 2004) was used at 1:8000 dilution in blocking buffer. For comparison between untagged NUF2 and NUF2-dTAG-FLAG protein levels antibody against NUF2 (abcam, Cat. No. ab176556) was used at 1:2000 dilution in blocking buffer.

### siRNA knockdown assay

siRNA knockdown of a select panel of genes identified in the screen was carried out by reverse transfecting siRNAs (Table S6, list of siRNAs used). Briefly, 150 nL of 5 µM siRNA stock solution was spotted on the 384 well glass bottom imaging plates using an Echo liquid handler following addition of 20 µL Optimem media (Thermo Fisher Scientific, Cat. No. 31-985-070) using a Multidrop dispenser. Required amount of Lipofectamine RNAiMAX (Thermo Fisher Scientific, Cat. No. 13778075) was added in each well using the Echo liquid handler and incubated for 30 minutes at room temperature to allow siRNA-lipofectamine complex to form. Next, the required number of cells diluted in 20 µLOptimem media containing 20% FBS (Gibco, Cat. No. 10082147) was dispensed into each well. The imaging plate was incubated for 72 hours at 37 °C in a cell culture incubator. Scrambled (Thermo Fisher Scientific, Cat. No. 4390846) and all-star cell death (siDEATH) (Qiagen, Cat. No. 1027298) siRNAs were used as controls to optimize the amount of lipofectamine RNAiMax reagent to achieve maximum transfection efficiency with minimum cytotoxicity.

### Statistical analyses

Statistical analysis including ANOVA, Tukey HSD test, and t-test with Bonferroni and FDR correction was performed using ggpubr (https://github.com/kassambara/ggpubr) and rstatix (https://github.com/kassambara/rstatix/releases) packages in R version 4.3.2. Statistical significance of difference was denoted by stars where * indicates p ≤ 0.05, ** indicates p ≤ 0.01, *** indicates p ≤ 0.001 and **** indicates p < 0.0001.

## Data Availability

Original image files used in this article are publicly available in figshare https://figshare.com/projects/Orderly_mitosis_shapes_interphase_genome_architecture/ 271192 and analysis code used for identification of hits in the CRISPR screens is available at GitHub: https://github.com/CBIIT/mistelilab-centromeres.

## Acknowledgements

We thank members of the Misteli lab for feedback throughout this project.This research was funded by the Intramural Research Program of the NIH, NCI, Center for Cancer Research through grant 1-ZIA-BC010309-25 to T.M and grant 1-ZIC-BC-011567 to HiTIF and in part with Federal funds from the National Cancer Institute, National Institute of Health under Contract No. HHSN26120150003I. Confocal imaging was performed in the CCR/ LRBGE Optical Microscopy Core, funded by the Intramural Research Program of the National Cancer Institute (NCI), Center for Cancer Research (CCR): project number ZIC BC 011574, and supported by Dr. T. S. Karpova. Fig. 6 was generated using BioRender. The content of this publication does not necessarily reflect the views or policies of the Department of Health and Human Services, nor does mention of trade names, commercial products, or organizations imply endorsement by the U.S. Government.

## Supplementary figures

**Fig. S1:**
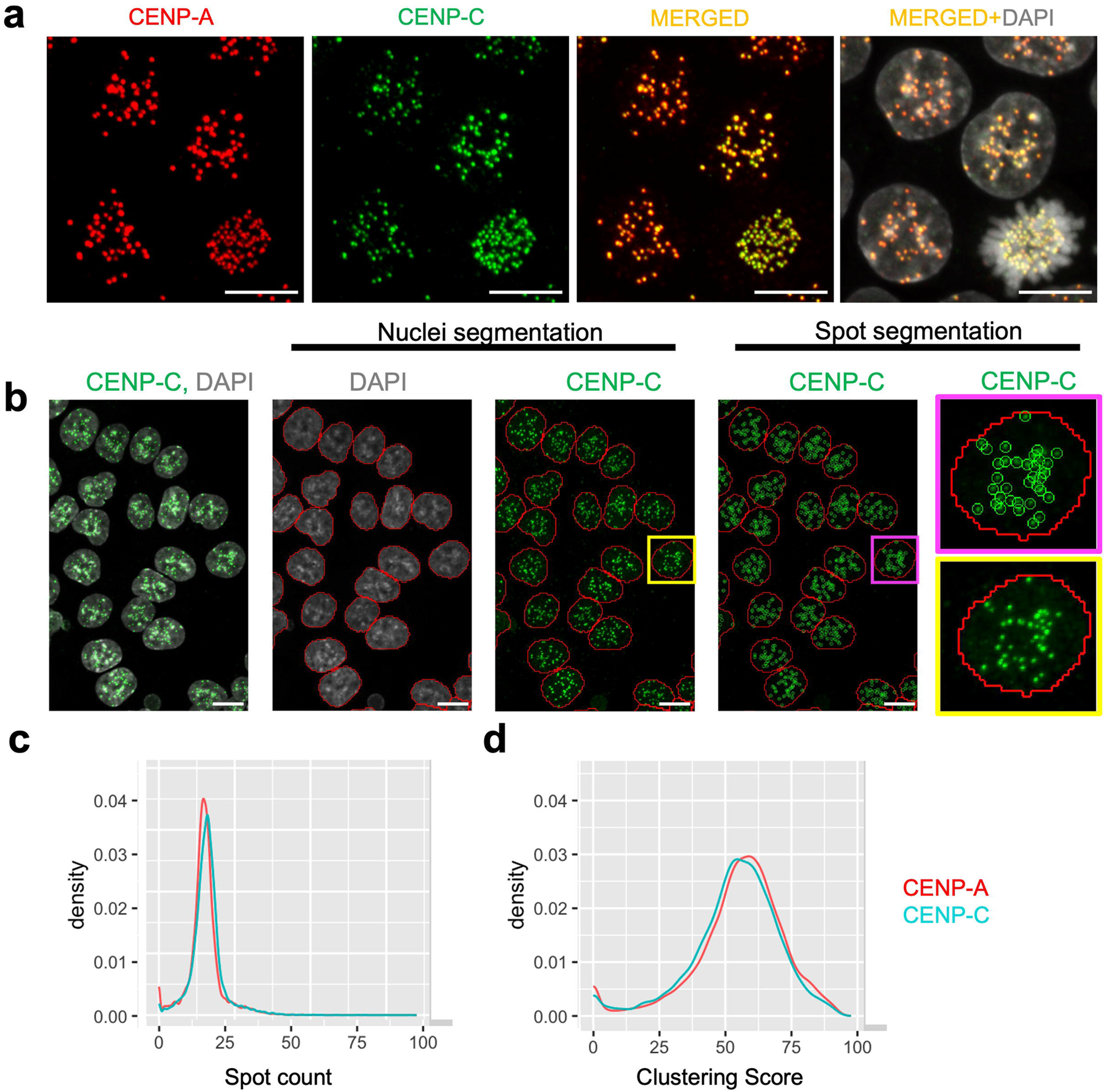
Quantification of centromere clustering using CENP-A and CENP-C as centromere markers. a, Co-staining of HCT116 cells with CENP-A (red), CENP-C (green) and DAPI (gray). Scale bar: 10 µm **b**, Representative image showing segmentation of DAPI stained nuclei (gray) and CENP-C-stained centromere spots (green) in high-throughput imaging data using HiTIPS. Red lines around the DAPI stained nuclei indicate nuclear segmentation, the green circles around CENP-C spots indicate segmentation of centromeres. Zoomed images of the same nucleus are shown with yellow and pink borders indicate before and after spot segmentation was applied. Scale bar: 10 µm. **c**, Quantification of spot count and **d**, Clustering Score using CENP-A (red) and CENP-C (blue) as centromere markers in HCT116 cells. Values are from two replicates, with at least 2000 cells analyzed for each experimental condition.

**Fig. S2.**
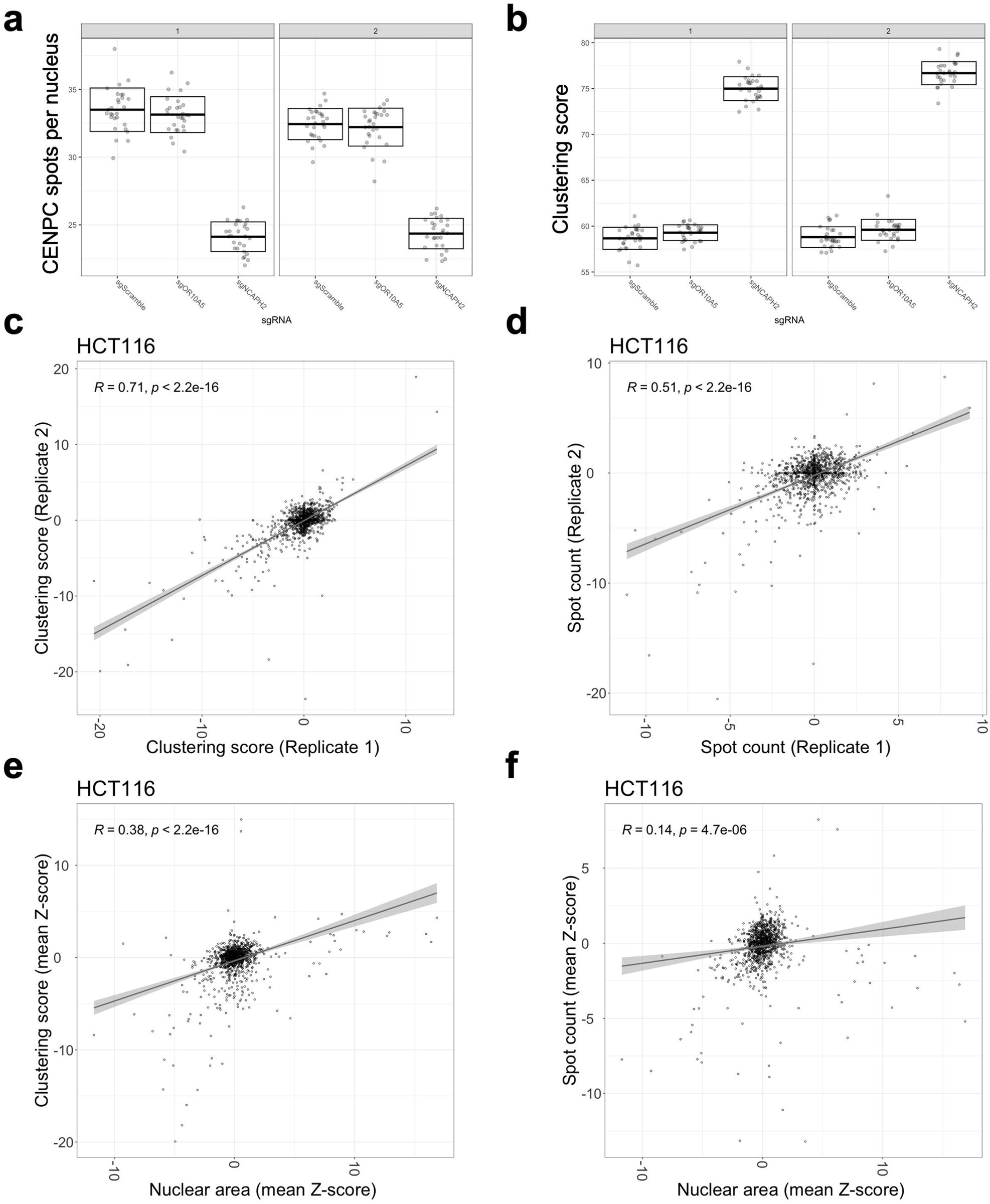
CRISPR knockout screens for centromere distribution phenotypes in HCT116 cells are reproducible. **a**, Mean and standard deviation for phenotypic separation between control sgRNAs with individual data points representing mean value per well for number of spots per nucleus and **b**, Clustering Score in two biological replicates. **c**, scatter plot showing changes in Clustering Score or **d**, spot count for replicate 1 (x-axis) and replicate 2 (y-axis) in HCT116 cells for each of the 1068 sgRNAs. A linear regression line (gray) was fitted to the data and Pearson’s correlation coefficient calculated is indicated at the top left corner of the plot. **e**, scatter plot and linear regression line correlating changes in Clustering Score (y-axis) and **f,** spot count (y-axis) with nuclear area (x-axis). Pearson’s correlation coefficients and corresponding p-values are indicated at the top left corner. Values are from 2 biological replicates. Typically, 200 to 500 cells were imaged for each target gene per experiment.

**Fig. S3:**
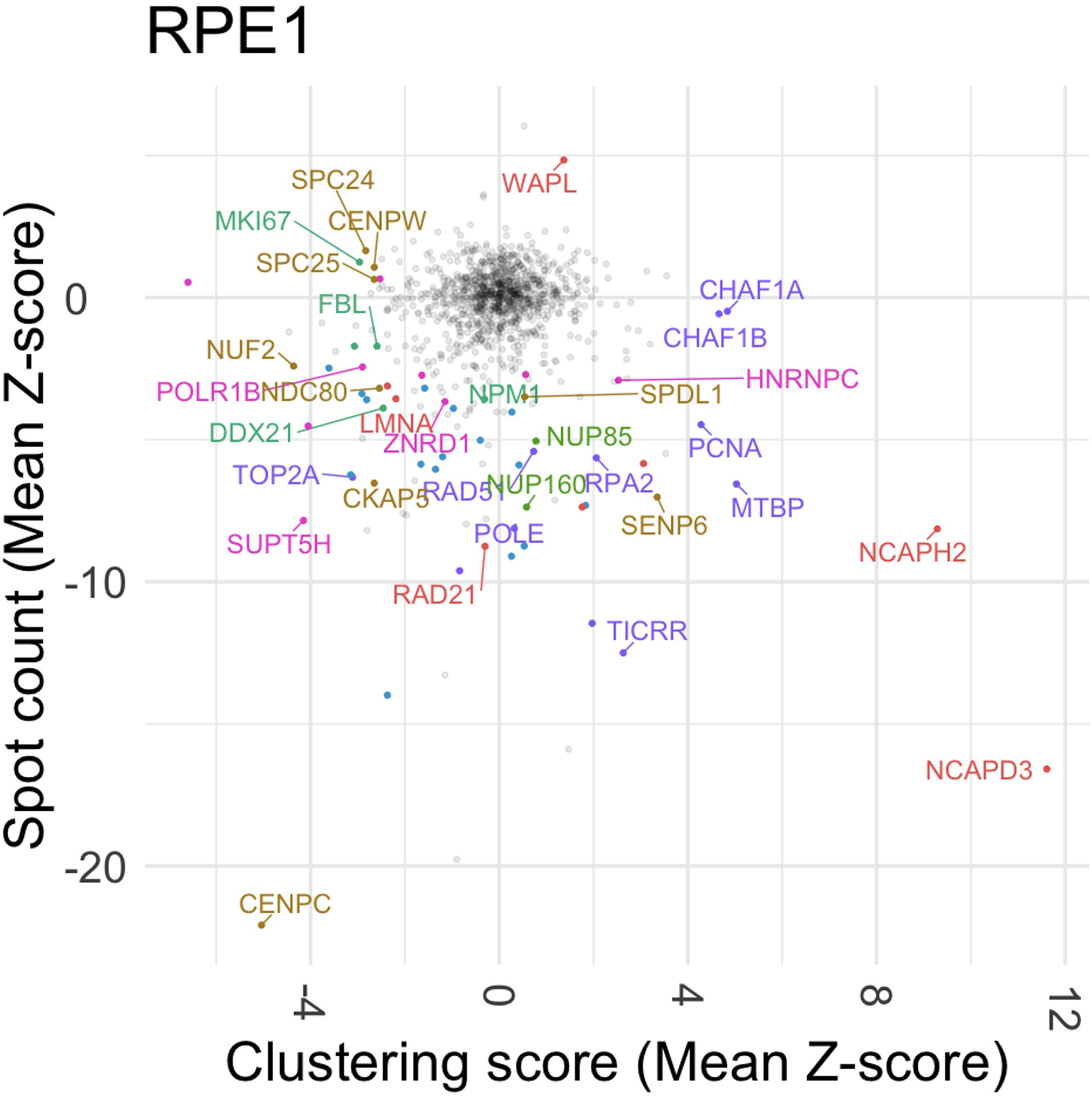
Identification of the molecular determinants of spatial centromere distribution in RPE1 cells. Changes in spot count (mean Z-score of two replicates, y-axis) and Clustering Score (mean Z-score of two replicates, x-axis) for each of the 1068 sgRNAs. The most prominent hits were labelled and color coded as in Fig. 2d. Non-hits are colored in gray. Values are from two biological replicates. Typically, 200 to 500 cells were analyzed for each target gene per experiment.

**Figure S4:**
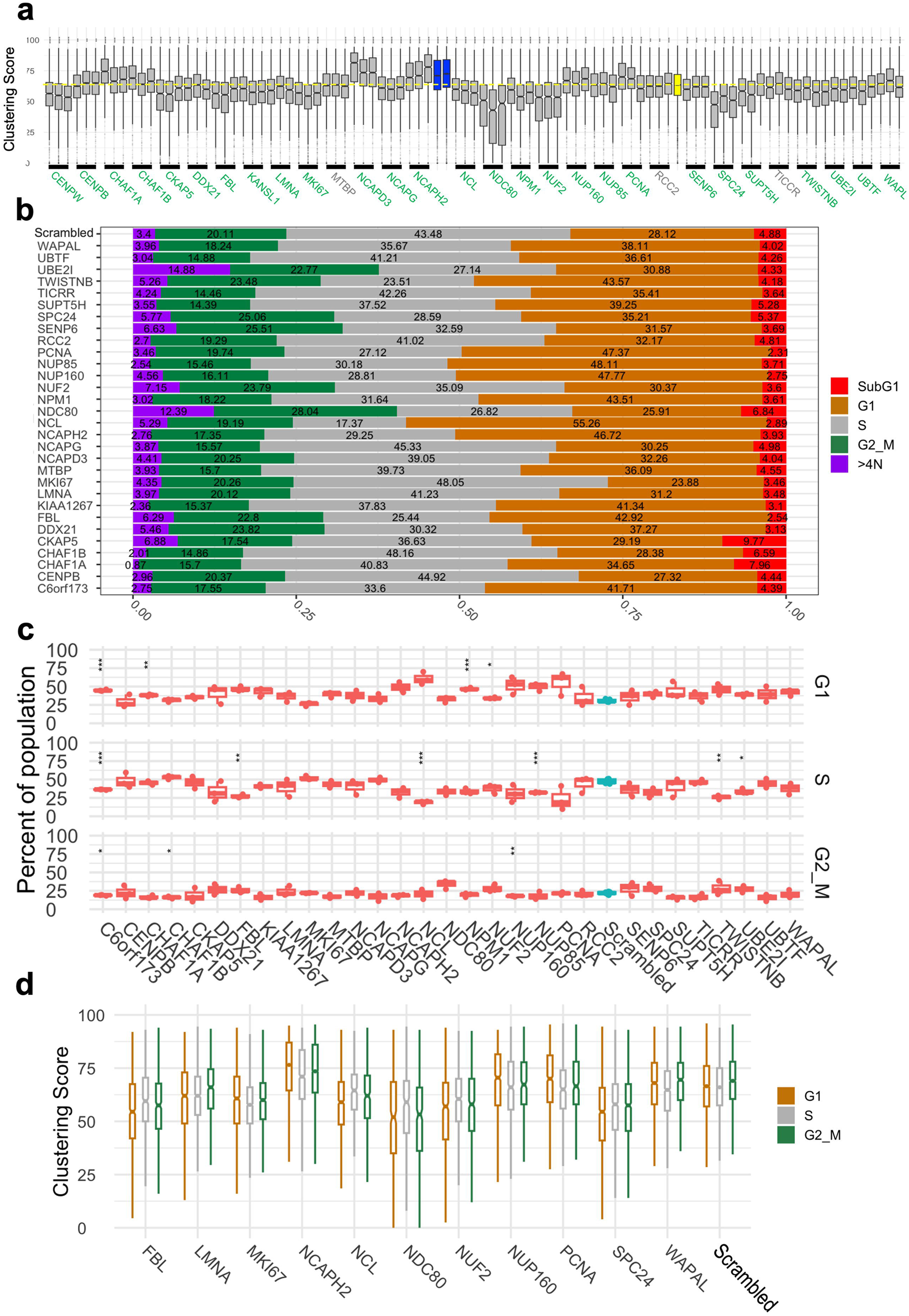
Validation of screen hits, cell cycle analysis of clustering factor knockdown and their effect on Clustering Score. **a**, Clustering scores of targets (x-axis) after siRNA knockdown in HCT116 cells. Two control siRNAs for siNCAPH2 are in blue and siScrambled is shown in yellow. Upon siRNA knockdown, the Clustering Score or spot count for the targets (x-axis) labelled in green change in the same direction as in the CRISPR-KO screens and are compared to the mean value for siScrambled as depicted by a horizontal yellow dotted line by performing pairwise t-tests with Bonferroni correction. Significantly different pairs are labelled with stars, where * indicates p ≤ 0.05, ** indicates p ≤ 0.01, *** indicates p ≤ 0.001 and **** indicates p < 0.0001. Targets in gray disagree with either Clustering Score or spot count or both parameters compared to data in CRISPR-KO screens. Three separate siRNA were used per target. **b**. Fraction of cells in each cell cycle stage (x-axis) after knockdown of select targets as indicated (y-axis). Individual sub-populations of the cell cycle are color-coded as identified using DAPI and EdU fluorescence intensity measurement, and their percentages are indicated. **c**, Bar plots showing the fraction of G1, S, and G2/M cells (y-axis) after siRNA knockdown of select targets (red) and scrambled siRNA control (blue). Statistical significance of difference (p<0.05) was tested using t-test with FDR correction as compared to the scrambled control and significantly different targets are labelled as stars, where * indicates p ≤ 0.05, ** indicates p ≤ 0.01, *** indicates p ≤ 0.001. Higher number of stars indicate lower p-value. **d**, Clustering score (y-axis) for select targets (x-axis) at G1 (brown), S (gray) and G2/M (green) stages. Values are from one representative experiment. Typically, 200 to 500 cells were analyzed per gene per experiment.

**Fig. S5:**
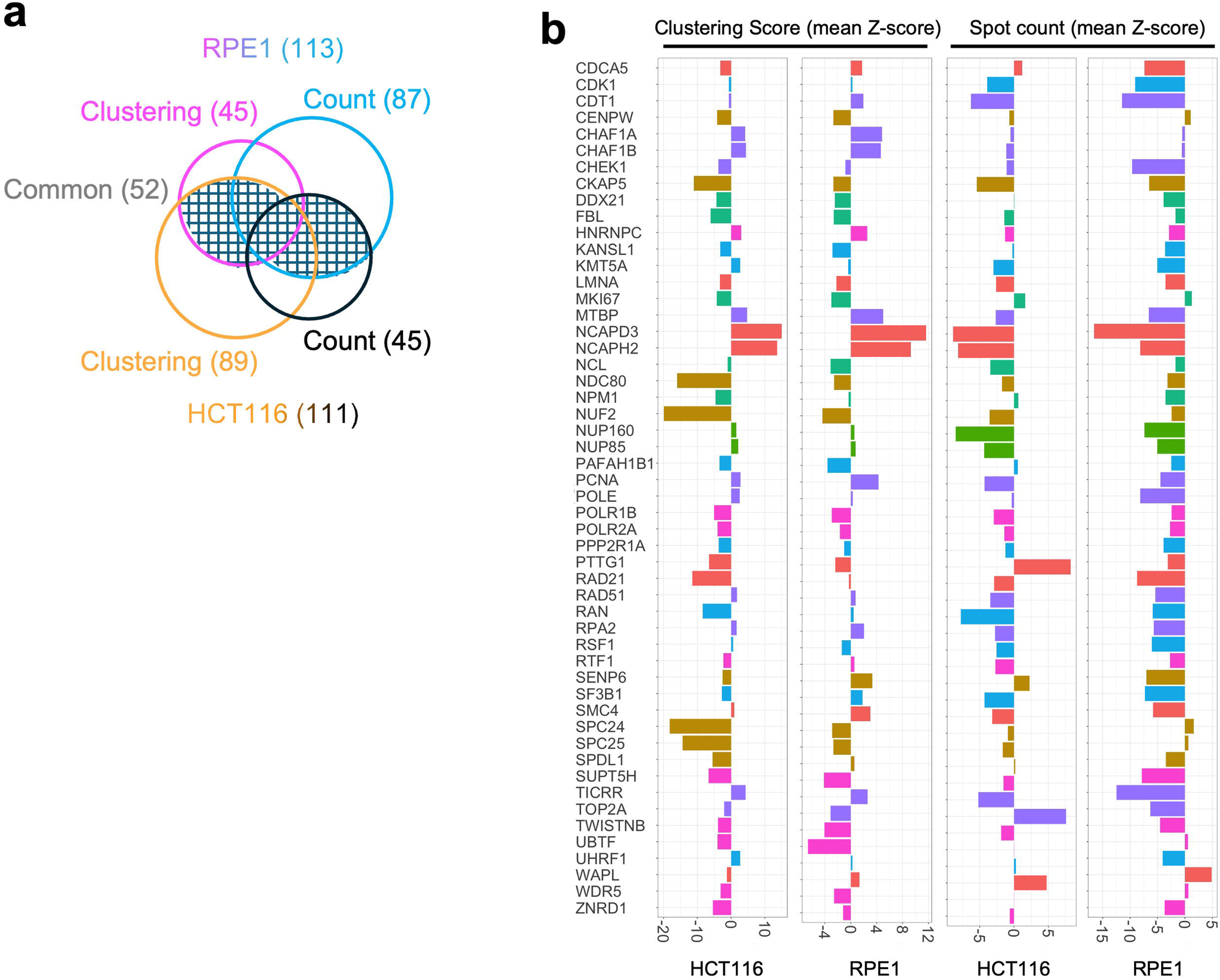
Comparative analysis of cell lines confirms common molecular determinants of spatial centromere distribution. a, 52 genes (black mesh) that are hits in both HCT116 and RPE1 cells. A total of 113 hits were selected in RPE1 cells for either Clustering Score (pink, 45) or spot count (blue, 87) and 111 hits in HCT116 cells for either Clustering Score (gold, 89) or spot count (black, 45). White non-shaded areas indicate unique hits in HCT116 (51) and RPE1 (53) cells. Values are from one representative experiment. Typically, 200 to 500 cells were analyzed for each target gene per experiment. **b**, Z-scores (x-axis) of spot count and Clustering Score for the 52 common hits (y-axis) in HCT116 and RPE1 cells. Genes are color coded based on their category as indicated in Fig. 2e.

**Figure S6:**
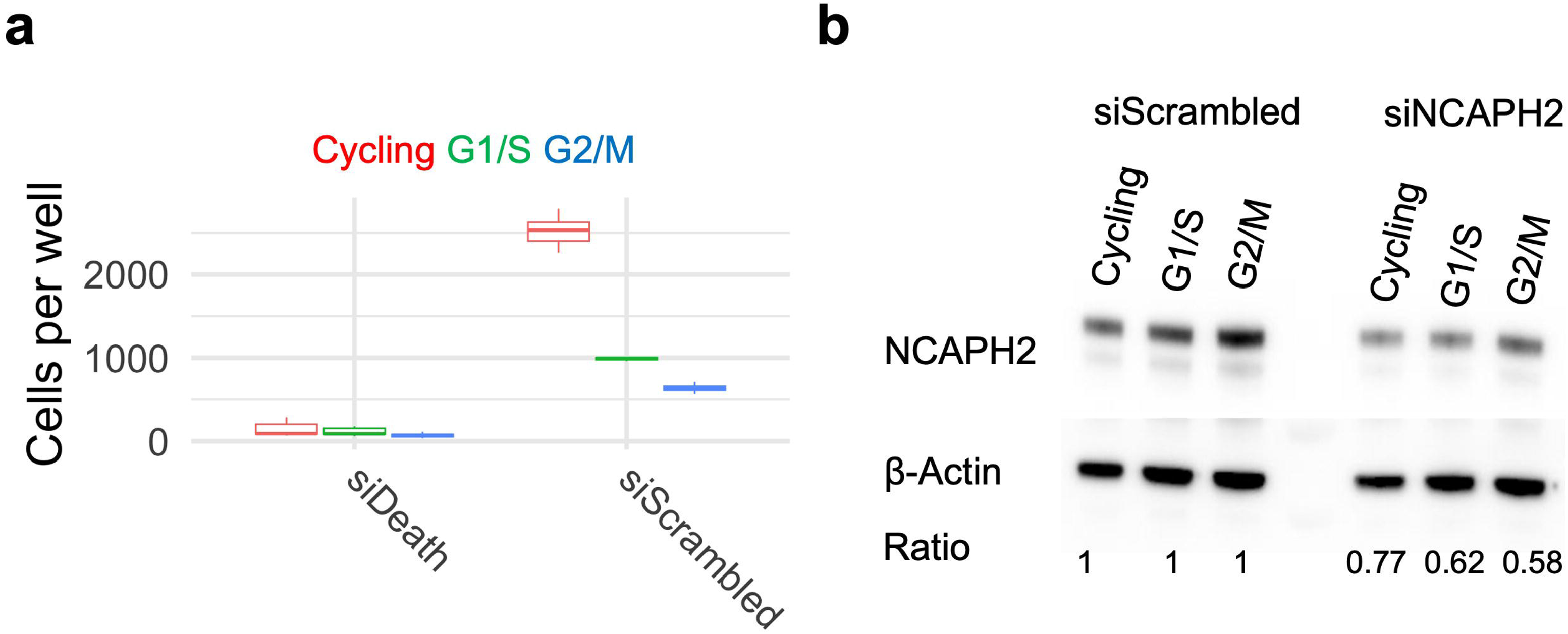
Validation of protein depletion in siRNA knockdown a, Number of cells per well at 72 hours after transfection of siDEATH and siScrambled control siRNAs in cycling (red), G1/S (green) or G2/M (blue) synchronized HCT116 cas9 cells as indicated. **b,** Western blots showing NCAPH2 protein levels after 72 hours of siRNA knockdown using siNCAPH2 and siScrambled in cycling, G1/S or G2/M synchronized HCT116 cas9 cells as indicated. NCAPH2 levels across samples were normalized using β-Actin and quantitated band intensities of NCAPH2 in siNCAPH2 samples were expressed as fraction of the corresponding siScrambled samples at cycling, G1/S and G2/M cells.

**Fig. S7:**
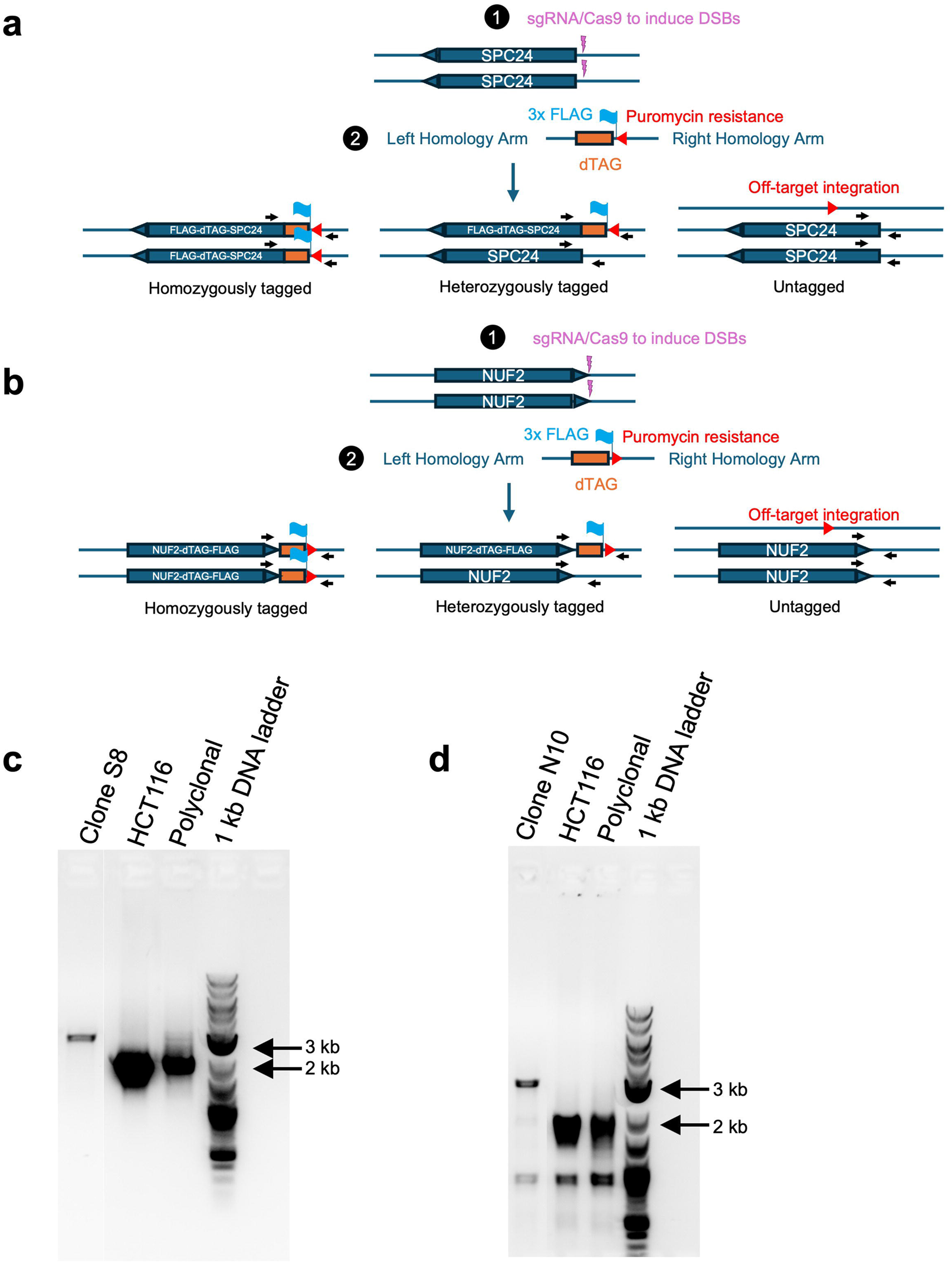
Construction and genotyping of FLAG-dTAG-SPC24 and NUF2-dTAG-FLAG cell lines. **a**, **b**, CRISPR knock-in strategy for homozygous tagging of SPC24 (**a**) and NUF2 (**b**) with the dTAG-FLAG epitope. Horizontal black arrows indicate positions of primers used for PCR confirmation of the tagged allele. **c**, **d**, PCR genotyping of single cell clone for FLAG-dTAG-SPC24 (**c**) and NUF2-dTAG-FLAG based on the strategy explained in **a** and **b**.

**Fig. S8:**
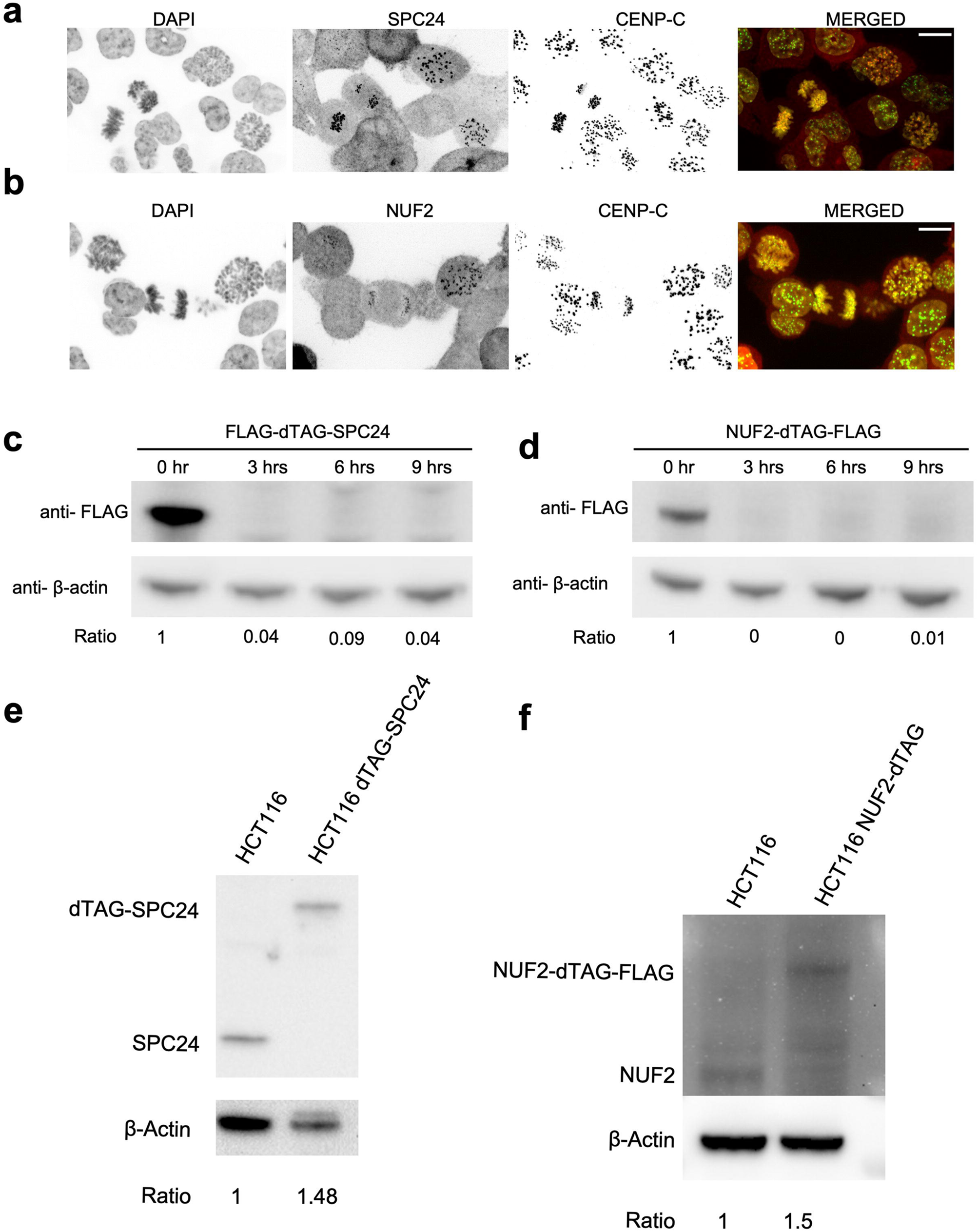
Characterization of FLAG-dTAG-SPC24 and NUF2-dTAG-FLAG cell lines. **a**, **b**, Representative images of knock-in cell lines expressing FLAG-dTAG-SPC24 (**a**), and NUF2-dTAG-FLAG (**b**) stained with DAPI (gray), CENP-C (green) and FLAG (red). Scale bar: 10 µm **c**, **d**, Westen blot images showing levels of FLAG-dTAG-SPC24 (**c**), and NUF2-dTAG-FLAG (**d**) at indicated time points after incubation with dTAG ligands and the relative ratios of dTAG-SPC24 or NUF2-dTAG to tubulin control compared to the level at the beginning of depletion (0 hr) are indicated below. **e**, **f**, western blots showing comparative levels of SPC24 and FLAG-dTAG-SPC24 (**e**) and NUF2 and NUF2-dTAG-FLAG (**f**) proteins in the indicated cell lines. Relative intensity ratio of FLAG-dTAG-SPC24 or NUF2-dTAG-FLAG proteins in the respective cell lines to the untagged SPC24 or NUF2 protein level in HCT116 cells is indicated.

**Fig. S9:**
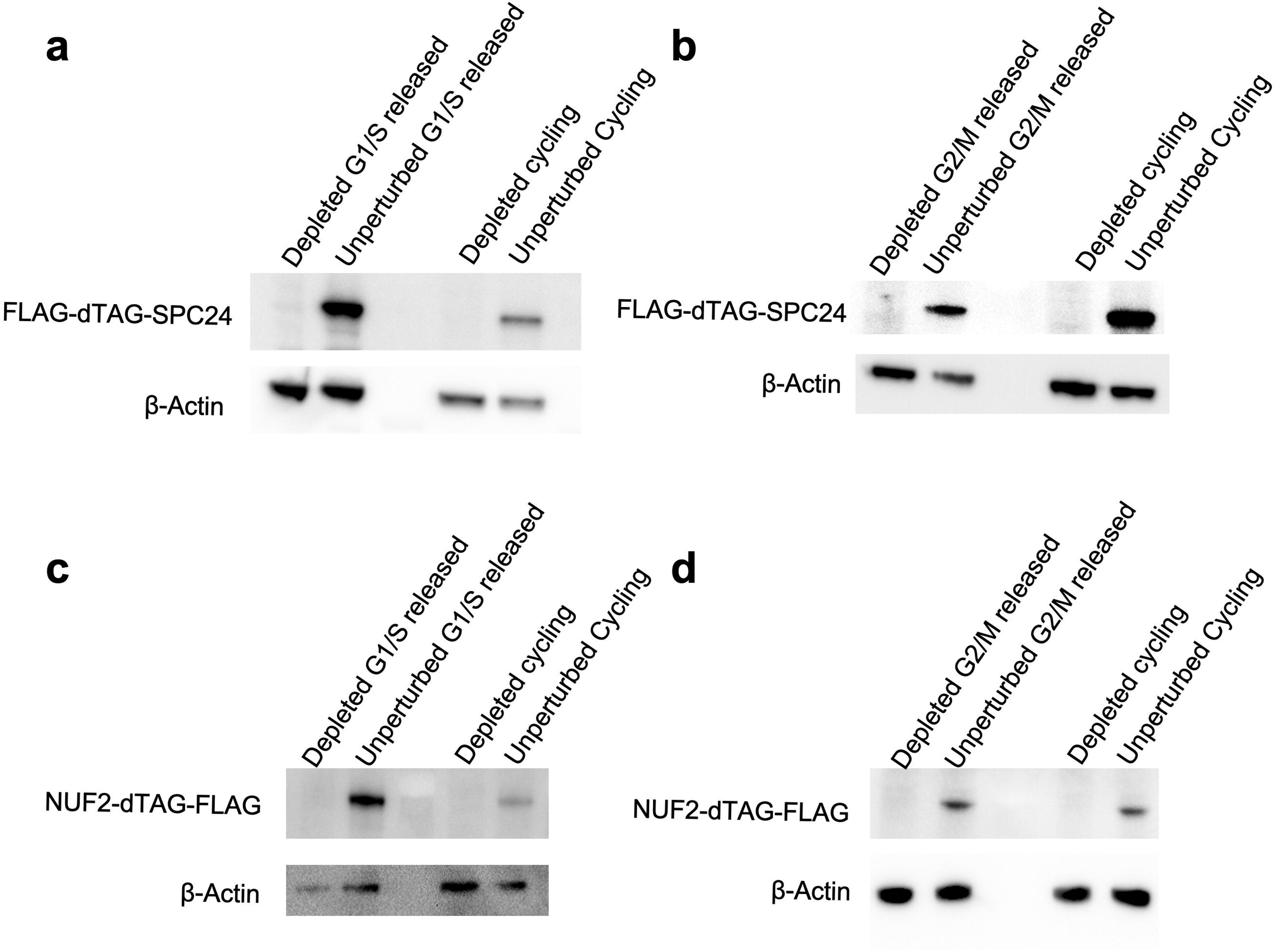
Quantification of the cell cycle stage specific depletion of FLAG-dTAG-SPC24 and NUF2-dTAG-FLAG. **a**, **b**, Western blots showing degron-based depletion of FLAG-dTAG-SPC24 in G1/S synchronized and cycling cells (**a**) and G2/M synchronized and cycling HCT116 cells (**b**). **c. d,** Western blots showing degron-based depletion of SPC24-dTAG-FLAG in G1/S synchronized and cycling cells (**c**) and G2/M synchronized and cycling HCT116 cells (**d**).

**Fig. S10:**
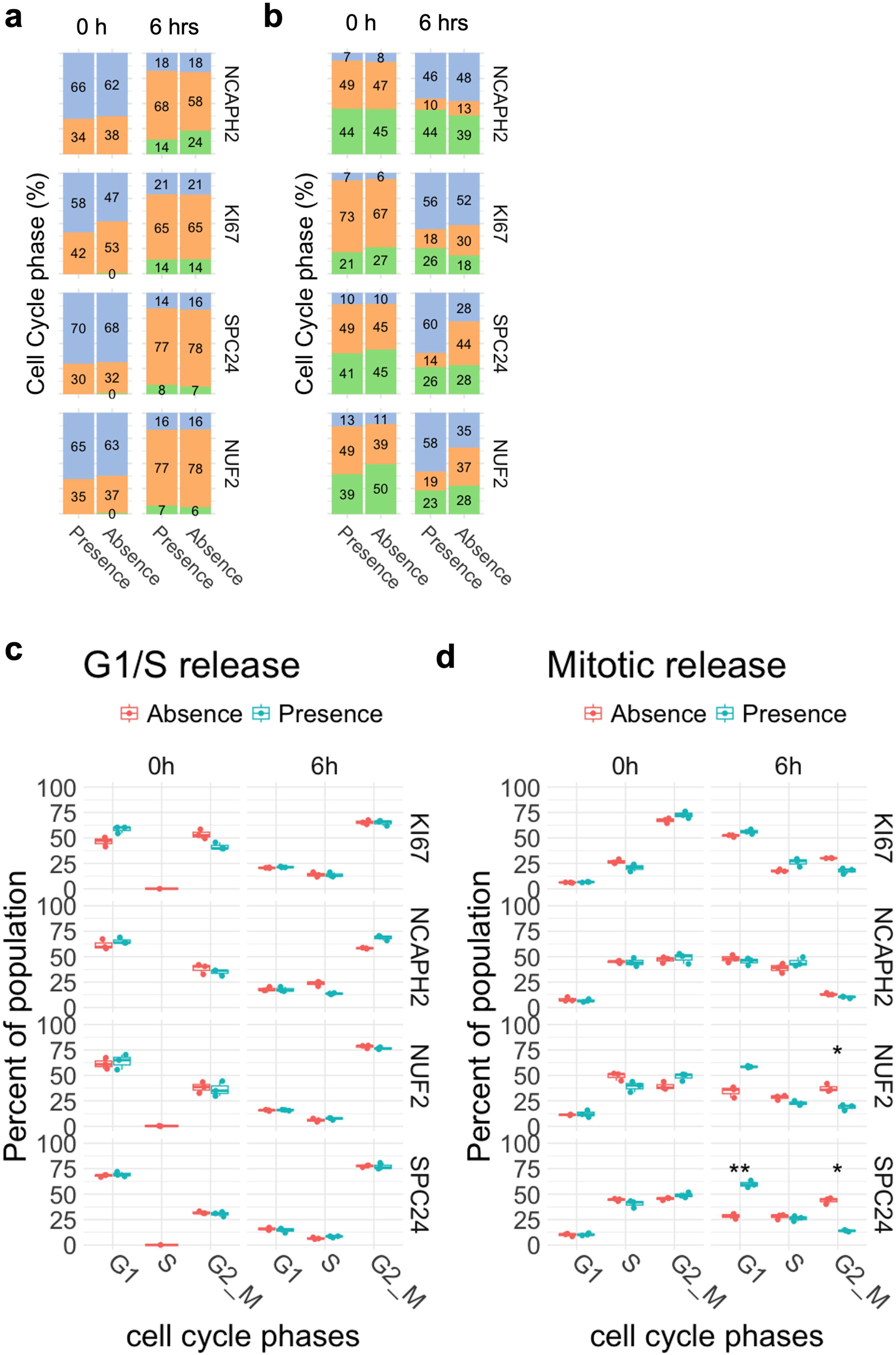
Quantification of cell cycle stages during G1 and mitotic release experiment. **a**, Fraction of G1 (blue), S (green) and G2/M (orange) cells were quantified before (0h) and after (6h) release from double thymidine block in the presence or absence of indicated clustering factors. **b**, Fraction of G1 (blue), S (green) and G2/M (orange) cells were quantified before (0h) and after (6h) release from G2/M block in the presence or absence of indicated clustering factors. Values are from one representative experiment containing three technical replicates. Typically, 200 to 500 cells were analyzed per sample. **c,** and **d,** shows percent of G1, S and G2/M cells (x-axis) in presence (blue) and absence (red) of indicated mitotic factors before (0h) and after (6h) release from G1/S block (**c**) and G2/M block (**d**). Statistical significance of difference was tested using t-test and significantly (p< 0.05) different pairs are indicated by stars, where * indicates p ≤ 0.05, ** indicates p ≤ 0.01. Higher number of stars indicate lower p-value.

**Fig S11:**
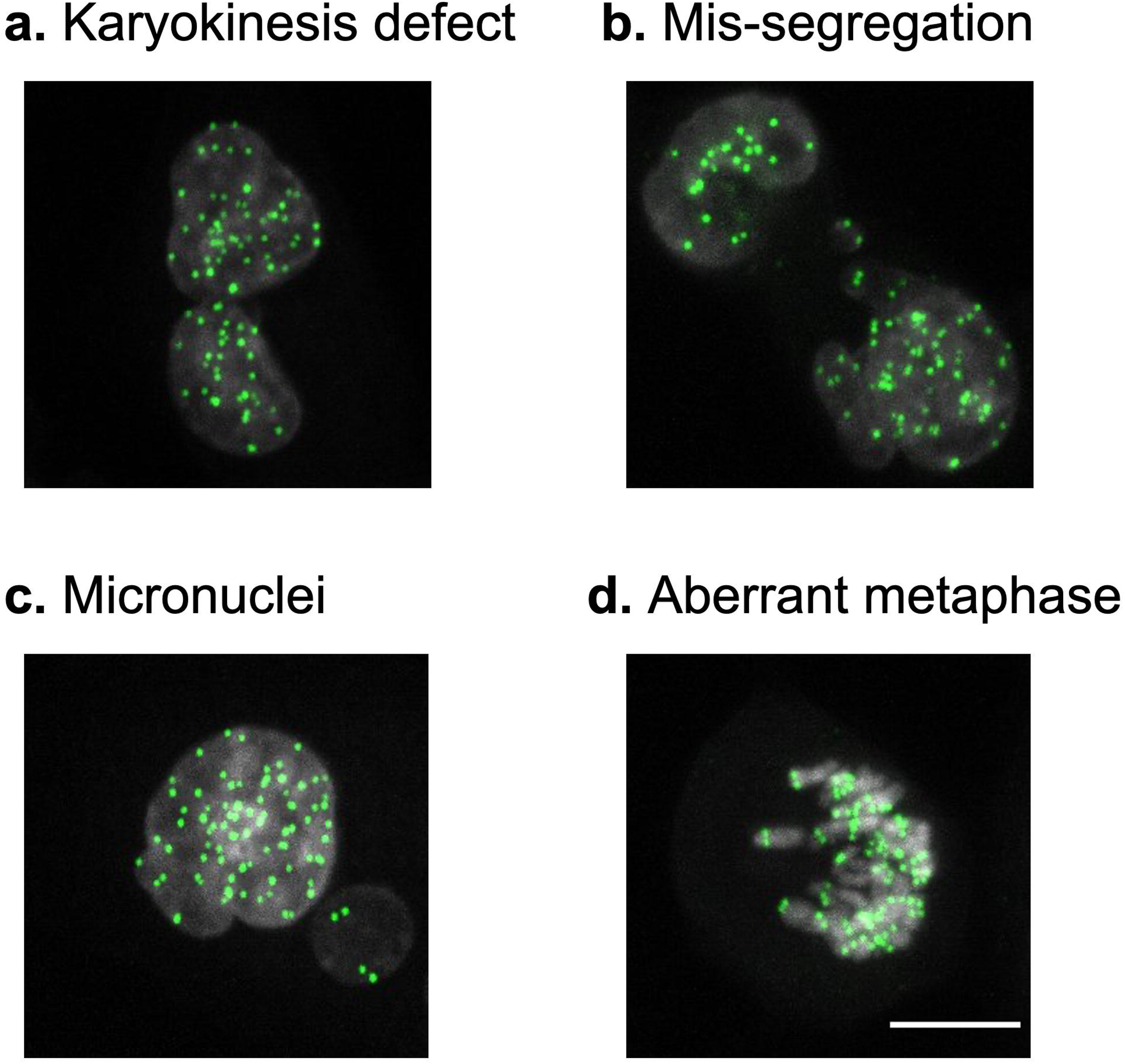
**Mitotic defects in the absence of SPC24. a-d**, Representative images showing examples of mitotic defects including karyokinesis defect (**a**), mis-segregation (**b**), micronuclei formation (**c**), and aberrant metaphase alignment (**d**) observed at 6 hours after G2/M arrested HCT116 cells were released in the absence of SPC24. Cells were immunofluorescent labelled with CENP-C (green) and DAPI (gray). Scale bar: 10 µm

